# The lncRNA *NEAT1_1* is dispensable for normal tissue homeostasis and cancer cell growth

**DOI:** 10.1101/574582

**Authors:** Carmen Adriaens, Florian Rambow, Greet Bervoets, Toomas Silla, Mari Mito, Tomoki Chiba, Asahara Hiroshi, Tetsuro Hirose, Shinichi Nakagawa, Torben Heick Jensen, Jean-Christophe Marine

## Abstract

*NEAT1* is one of the most studied lncRNAs, in part because its silencing in mice causes defects in mammary gland development and corpus luteum formation, and protects them from skin cancer development. Moreover, depleting *NEAT1* in established cancer cell lines reduces growth and sensitizes cells to DNA damaging agents. However, *NEAT1* produces two isoforms and because the short isoform, *NEAT1_1*, completely overlaps the 5’ part of the long *NEAT1_2 isoform*, the respective contributions of each of the isoforms to these phenotypes has remained unclear. Whereas *NEAT1_1* is highly expressed in most tissues, *NEAT1_2* is the central architectural component of paraspeckles, which are nuclear bodies that assemble in specific tissues and cells exposed to various forms of stress. Using dual RNA-FISH to detect both *NEAT1_1* outside of paraspeckles and *NEAT1_2/NEAT1* inside this nuclear body, we report herein that *NEAT1_1* levels are dynamically regulated during the cell cycle and targeted for degradation by the nuclear RNA exosome. Unexpectedly, however, cancer cells engineered to lack *NEAT1_1*, but not *NEAT1_2*, do not exhibit cell cycle defects. Moreover, *Neat1_1*-specific knockout mice do not exhibit the phenotypes observed in *Neat1*-deficient mice. We propose that *NEAT1* functions are mainly, if not exclusively, attributable to *NEAT1_2* and, by extension, to paraspeckles.

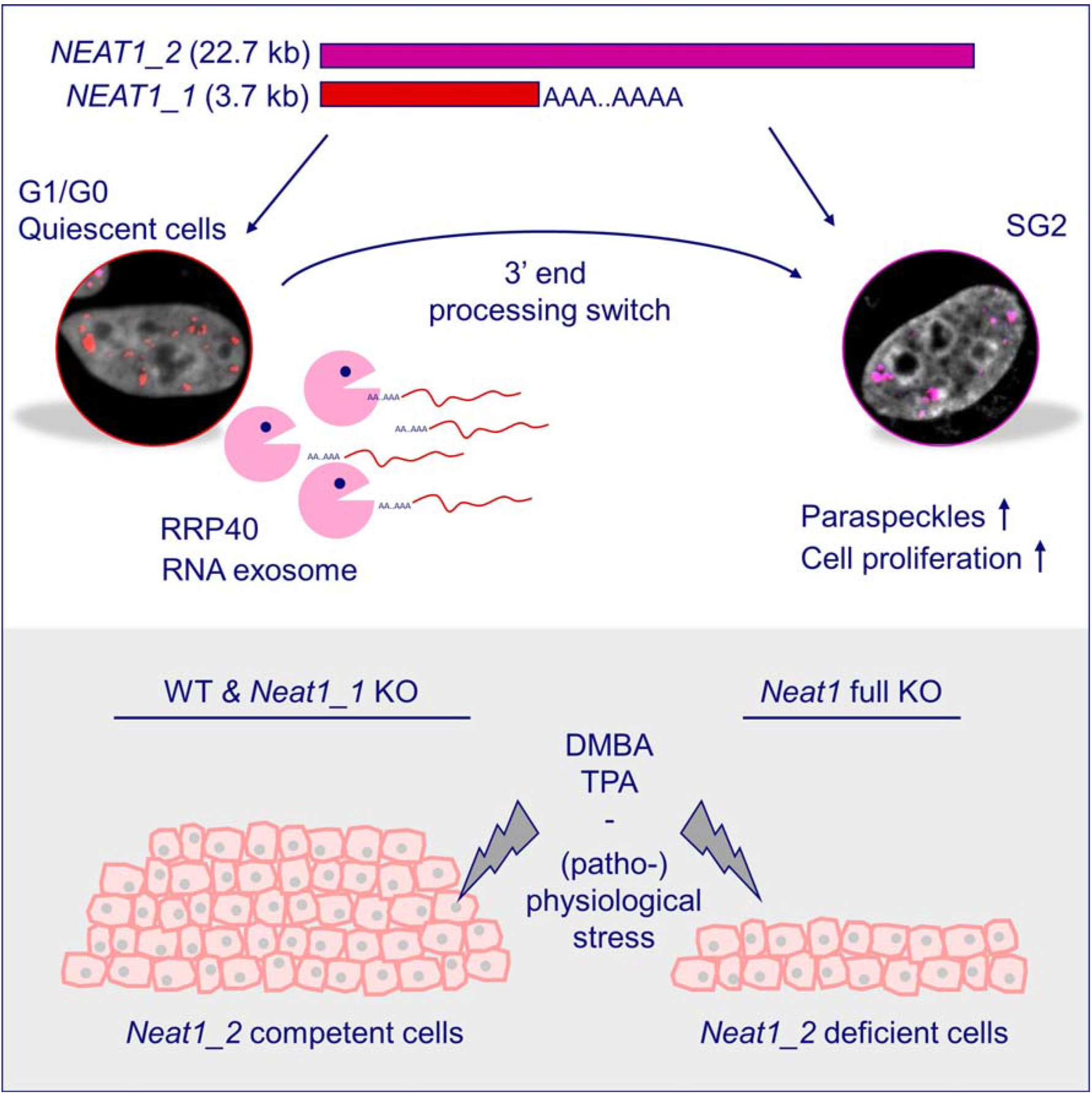
Graphical abstract.

## Introduction

Long non-coding RNAs (lncRNAs) exceed 200 nucleotides in length and lack protein coding potential. In the past decade, some of these molecules have arisen as prominent players in a range of cellular processes, including the formation of gene regulatory domains, the spatial organization of the genome or cell plasticity (Quinn and Chang 2016). One of these lncRNAs, *NEAT1*, is required for the assembly of intriguing and enigmatic nuclear bodies known as paraspeckles (PS) (Sunwoo et al. 2009; Sasaki et al. 2009; Clemson et al. 2009). Since then, PS have been implicated in gene expression regulation and in the maintenance of DNA integrity in response to endogenous and exogenous forms of stresses (Chen and Carmichael 2009; Hirose et al. 2014; Imamura et al. 2014; Choudhry et al. 2014; Adriaens et al. 2016; Mello et al. 2017; Ahmed et al. 2018; Lellahi et al. 2018). This may occur through their interaction with the RNA interference (RNAi) machinery and with micro-RNAs (Jiang et al. 2017; Li et al. 2018; Shuaib et al. 2019), or via the modulation of transcriptional and post-transcriptional regulators (Hirose et al. 2014; Hu et al. 2015; Kawaguchi et al. 2015; Torres et al. 2016; Morchikh et al. 2017; Wang et al. 2018; Hupalowska et al. 2018). Importantly, the generation of *Neat1*-deficient mice has highlighted a critical role for this lncRNA in the formation of a functional lactating mammary gland and corpus luteum (Nakagawa et al. 2014; Standaert et al. 2014; Adriaens et al. 2016). *NEAT1* was also shown to protect pre-neoplastic cells from accumulating excessive DNA damage and, thereby, to be required for tumor initiation (Adriaens et al. 2016).

Because of the above, *NEAT1* has emerged as one of the most studied lncRNAs. However, several outstanding questions remain regarding NEAT1 biology. The *NEAT1* locus produces two different lncRNAs: a long transcript of ∼22.7 kb, *NEAT1_2*, and *NEAT1_1*, a shorter transcript of ∼3.7 kb (Sasaki et al. 2009). One of the key unanswered questions to date is what the actual contributions of these two distinct isoforms are to the above-described phenotypes.

*NEAT1_1* is a highly conserved and abundant poly-adenylated transcript, which is detected in virtually all tissues (Nakagawa et al. 2011). In contrast, expression of *NEAT1_2*, which is required for PS assembly, is only detected under specific physiological conditions (i.e. mammary gland development, corpus luteum formation) and in response to various forms of stress, including oncogenic stress. Conversely, non-differentiated cells preferentially produce *NEAT1_1*, and consequently lack *NEAT1_2* and thereby PS (Sunwoo et al. 2009; Chen and Carmichael 2009; Modic et al. 2018). Interestingly, PS appear in over 65% of human epithelial cancers (Adriaens et al. 2016), where they predict poor prognosis (Li et al. 2018). In contrast, they are either completely absent or only sporadically detected in the adjacent normal tissues (Adriaens et al. 2016).

*NEAT1_2* is a read-through transcript that is produced as a result of incomplete processing of the 3’-end of *NEAT1_1*. Little is known about the mechanisms that regulate *NEAT1_1* 3’-end processing, other than that it depends on the activity of an ubiquitous nucleic acid binding protein, hnRNP K, and the 3’-end cleavage factor Im (CFIm) complex (Naganuma et al., 2012). PS assembly therefore depends on this poorly understood switch from transcriptional termination to read through (Naganuma et al. 2012; Yamazaki et al. 2018). Because PS are detected in the cellular compartments that exhibit phenotypes following silencing of the *Neat1* locus, it has been tempting to speculate that these defects arose as a consequence of loss of *Neat1_2* and PS. However, the investigated mice were also deficient for *Neat1_1* and its contribution to these phenotypes has therefore remained unclear. The complete overlap of *NEAT1_1* with the 5’ end of NEAT1_2 factors has made it particularly challenging to study the individual contribution and behavior of these two isoforms independently. As a result, most groups that study NEAT1 biology do not discriminate whether the observed effects in *NEAT1* perturbation experiments are attributable to *NEAT1_1*, *NEAT1_2* or both.

In order to study whether the two isoforms functionally interact, as recently proposed (Fox et al. 2018), or whether they exert distinct biological functions, we employed dual RNA-FISH, isoform specific gene editing and knockdown strategies. We show that the two isoforms are differentially expressed at various phases of the cell cycle and that *NEAT1_1* is a target for degradation by the nuclear RNA exosome machinery. However, despite the high evolutionary conservation, the ubiquitous expression and its tight regulation between the cell-cycle, mice and cells deficient for *NEAT1_1* did not exhibit any of the phenotypes observed upon ablation of both isoforms or *NEAT1_2* only. Moreover, the phenotypes observed upon silencing *NEAT1_2* in *NEAT1_1*-proficient cells were recapitulated in *NEAT1_1*-deficient cells. We propose that *NEAT1*’s biological functions are solely attributable to the *NEAT1_2* isoform, and by extension to PS formation. The pathophysiological function of *NEAT1_1*, if any, remains to be elucidated. Our study therefore encourages a more careful dissection of individual non-coding RNA isoforms and indicates that high abundance and conservation is not necessarily predictive of functionality.

## Results

### Differential regulation of *NEAT1* isoforms in response to stress

To dissect a putative differential behavior of the two *NEAT1* isoforms in cultured cancer cells, we performed RNA-FISH with two distinct probes that target both transcripts (red), or *NEAT1_2* specifically (blue) (Figure 1A). As the first portion of NEAT1_2 completely overlaps the short isoform, a pink signal (red+blue) marks the presence of both transcripts, whereas red signals indicate the sole presence of *NEAT1_1*, outside of PS. Note that this approach does not allow to determine whether *NEAT1_1* localizes to PS (Clemson et al. 2009; Souquere et al. 2010). Using this method, we observed a fraction of untreated, proliferating U2OS cells displaying *NEAT1_1* in the nucleoplasm, outside of PS [37.7 ± 15.8 % of the cells] (Figure 1B, C, left panel and box plot). U2OS cells are triploid for *NEAT1* and, consistently, often three pink dots were detectable, indicating PS formation at those loci (Clemson et al. 2009; Mao et al. 2011). We and others have shown that induction of p53 stimulates transcription of *NEAT1* and PS formation (Blume et al. 2015; Adriaens et al. 2016; Idogawa et al. 2017; Mello et al. 2017). Accordingly, treatment of the cells with the p53 inducer Nutlin-3a increased the size of PS. This was accompanied by a dramatic increase in the proportion of cells displaying nucleoplasmic *NEAT1_1*-specific signal (79.0 ± 8.3 % of the cells; Figure 1B, C center panel and box plot). In contrast, exposure to the DNA damaging agent Hydroxyurea (HU), decreased the *NEAT1_1* specific signal (with only 5.2 ± 3.7 % of the cells being positive; Figure 1B, C, right panel and box plot).

**Figure 1:**
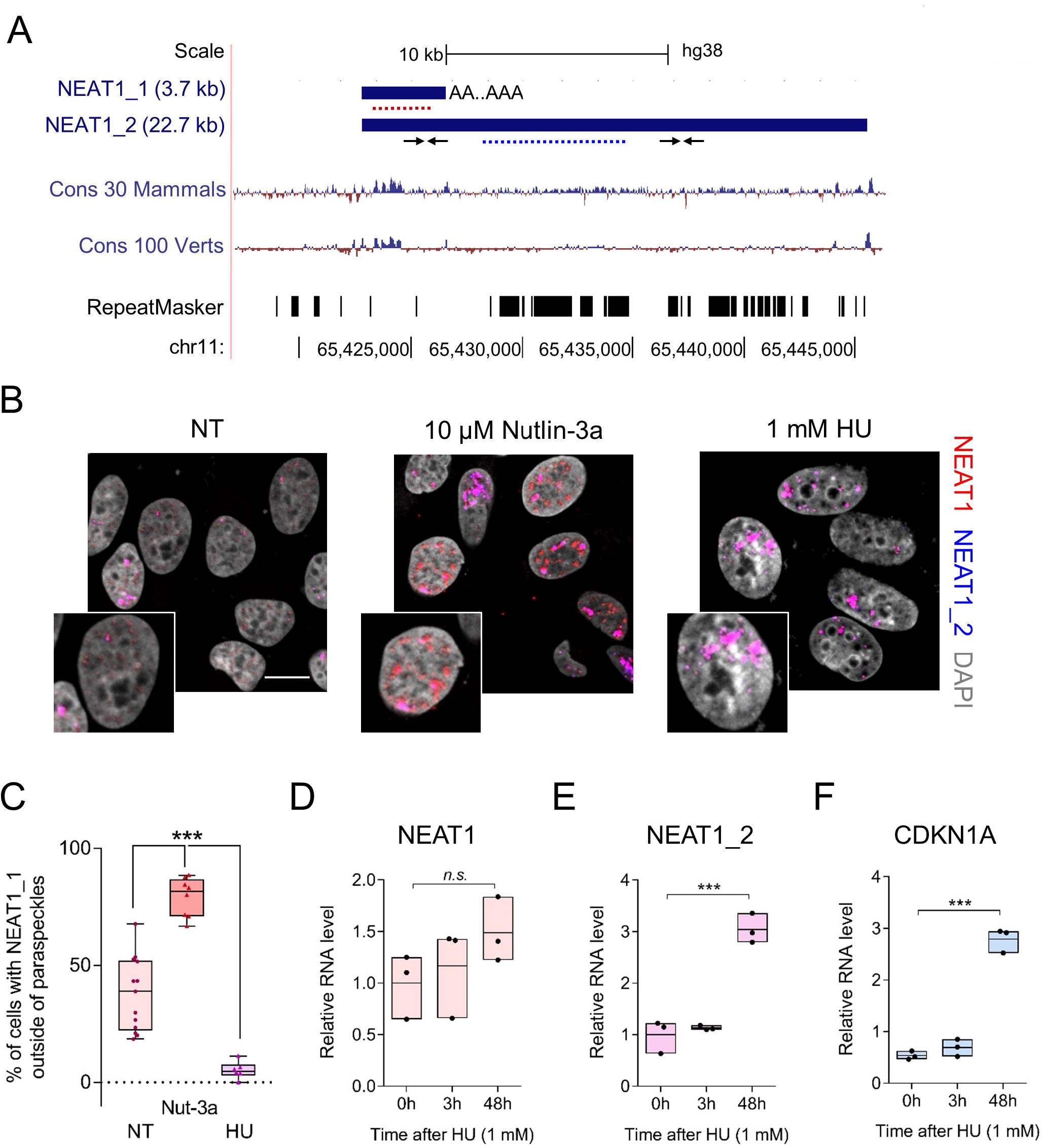
*NEAT1* isoforms are differentially regulated by distinct p53 activating agents. A. Refseq representation of human *NEAT1* isoforms in the UCSC genome browser (hg38); nucleotide-level conservation among 30 mammals and 100 vertebrates and the location of repeats in the genome sequence. Red and blue dotted lines represent RNA-FISH probes targeting both and the long *NEAT1_2* isoform specifically, respectively. Note that when targeting the long isoform, blue and red probes will overlap and thus show paraspeckles in pink. Small arrows represent approximate locations of RT-qPCR primers used in this study. B. Representative confocal images of RNA-FISH targeting *NEAT1* isoforms in U2OS cells in non-treated (NT), 10 µM Nutlin-3a (24h) and 1mM hydroxyurea (HU, 48h) conditions. Scale bar, 15 µm. C. Quantification of the percentage of cells in which the short isoform can be observed outside of paraspeckles. Each dot represents an independent experiment (N = 13, 8, and 6, respectively). D, E and F. Relative levels of *NEAT1* (both isoforms, D), *NEAT1_2* (E) and the canonical p53 target *CDKN1A* (F) after 0, 3 and 48h of HU treatment (1mM).

Real time quantitative PCR (RT-qPCR) analyses with primers detecting both isoforms and *NEAT1_2* only (Figure 1A for primer locations) established that *NEAT1_2* was specifically upregulated in cells exposed to HU (Figure 1D, E). As expected, an increase in the levels of the p53-target *CDKN1A* was also observed, indicating its transcriptional activation (Figure 1F). Although we noted that the sizes of *NEAT1_2*-containing bodies slightly decreased in these cells, we confirmed that they were genuine PS by co-staining with NONO, a canonical PS marker (Fox et al. 2005; Souquere et al. 2010) (Figure S1).

These data indicated that the ratio and localization of *NEAT1_1* and *NEAT1_2* vary depending on the type of stress inflicted to the cells. The experiments also highlighted the presence of a large pool of *NEAT1_1* that does not overlap with *NEAT1_2*-containing PS (Nakagawa et al. 2011; Li et al. 2017).

### *NEAT1_1* levels are dynamically regulated during the cell cycle

In the p53 competent U2OS cells, Nutlin-3a induces primarily a G1 cell cycle block through activation of *CDKN1A* (Shen et al. 2008). In contrast, HU arrests cells in S phase through the inhibition of the deoxynucleotide (dNTP) producing enzyme ribonucleotide reductase (RNR), thereby depleting the dNTP pool during replication (Singh and Xu 2016). We therefore hypothesized that *NEAT1_1* and *NEAT1_2* levels may be differentially regulated during the cell cycle. To test this, we deprived cells from serum to halt them in a resting, G0-like state (G0). Subsequently, cells were released in 20% serum in the presence of the DNA polymerase inhibitor Aphidicolin to synchronize them at the G1/S phase boundary. DNA content analysis using flow cytometry confirmed efficient synchronization of cells (Figure 2A). Using the above described dual RNA-FISH strategy, we observed that 87 ± 15% of G0 halted cells expressed the short *NEAT1_1* isoform outside of PS (Figure 2B, C; Figure S2A). In contrast, only 4.7 ± 7.8% of the G1/S arrested cells displayed *NEAT1_1*-specific signals, that did not overlap with *NEAT1_2* (Figure 2B, C; Figure S2A, S3A). RT-qPCR analysis demonstrated that total levels of *NEAT1*, but not of *NEAT1_2*, were increased in G0 cells (Figure S3C). Contrastingly, in G1/S cells *NEAT1* and *NEAT1_2* levels were comparable and significantly lower as compared to non-synchronized cells (*p* = 0.0018 for *NEAT1* and *p* = 0.0077 for *NEAT1_2*, unpaired two-sided t-test using Holm-Sidak method) (Figure S3D). Whereas G0 cells displayed on average three *NEAT1_2*-containing PS and 24 ± 23 *NEAT1_1* RNA FISH signals, respectively (Figure S2B, D), G1/S cells displayed on average 4.5 PS per cell (Figure S2C). Together, these data indicated that, in non-proliferating cells, *NEAT1_1* is the predominant isoform, and that, upon entering the cell cycle, the amount of signal for *NEAT1_*1 outside of PS abruptly drops.

**Figure 2:**
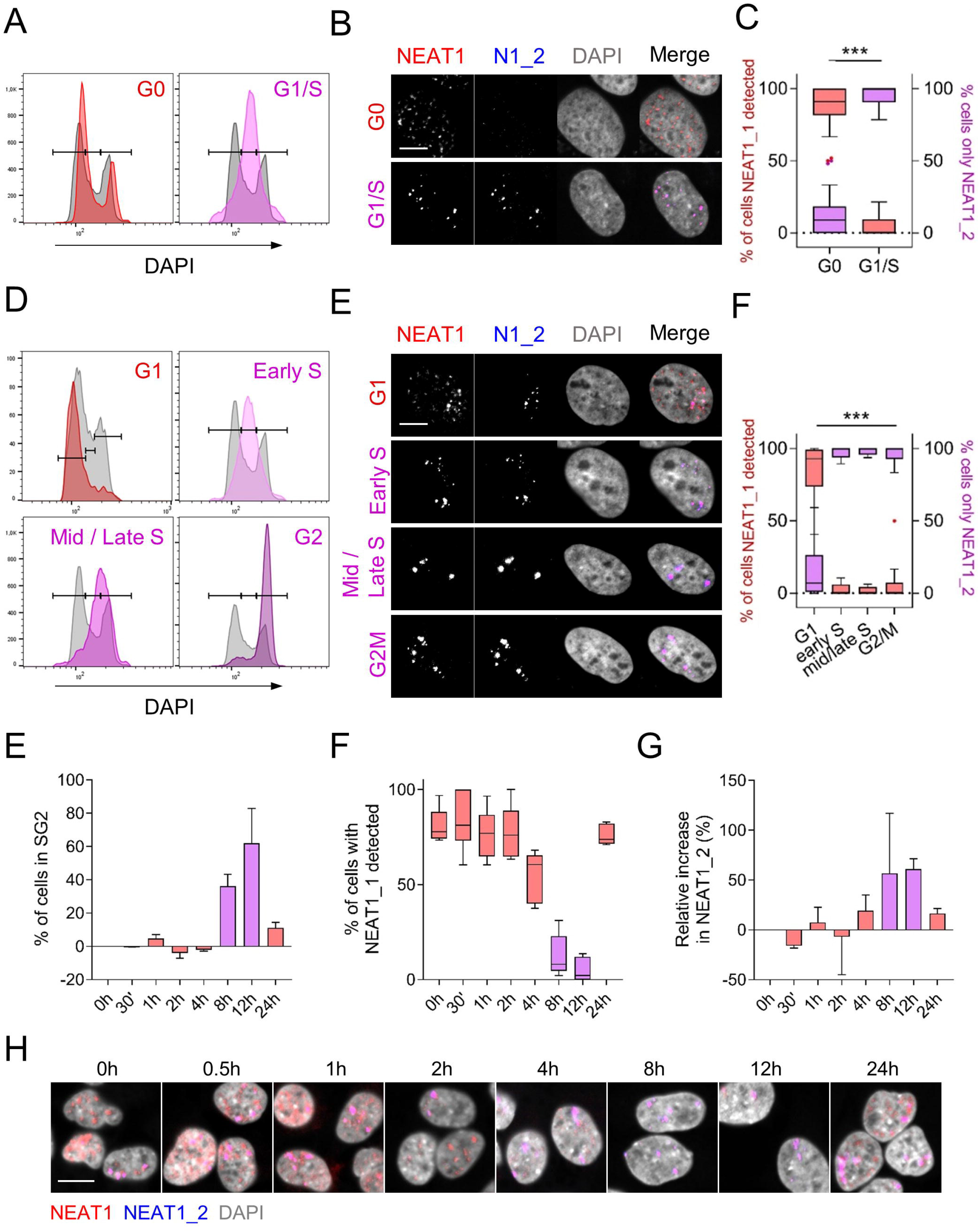
*NEAT1* isoforms are differentially regulated during cell cycle. A. Representative DNA content (DAPI) distribution of U2OS cells in G0 (3 days starvation) and G1/S (G0 cells released in 20% serum + 5 µg/ml Aphidicolin for 24h). Grey background plots are control non-synchronized (NS) cells in culture. B. Representative images of *NEAT1/NEAT1_2* RNA FISH of the cells in A. Scale bar, 10 µm. C. Quantification of the % of cells in which *NEAT1_1* was detected independently of *NEAT1_2* paraspeckles, red boxplots, left y-axis. Percentage of cells in which only *NEAT1_2*/paraspeckles were detected, right y-axis, purple boxplots. Tukey plots of individual data points (1 point per picture). Significance was calculated using an unpaired, two-sided t-test on independent biological replicates (N = at least three). ***, p<0.001. D. Like in A, but from U2OS cells synchronized by double thymidine block and released for 18 (G1), 2 (early S), 4 (mid/late S) and 8 hours. E. Representative images of *NEAT1/NEAT1_2* RNA FISH of the cells in D. F. Same as C for the cells in D. Significance was calculated using a 2-way ANOVA with Dunnett’s correction for multiple comparisons on independent replicates (N = at least three). ***, p<0.001 for G1 vs early S, mid/late S, and G2). N1_2 = *NEAT1_2*. E. % of cells in S and G2 phases at different time points relative to time = 0h in HeLa cells upon release into 20% serum after 3 days starvation. F. Box plots (Min to Max) of individual data points (N = 5 and 3 pictures per replicate, respectively) quantifying the % of cells in E in which *NEAT1_1* was detected using RNA-FISH against both and the long isoform specifically. G. Relative increase of *NEAT1_2* by RT-qPCR of the cells in E. Error bars are standard deviation of N = 2 experiments in E and G. H. Representative pictures of cells in E. Scale bar, 20 µm.

To substantiate these data, we used a conventional double thymidine block-and-release protocol to synchronize cells and subsequently release them for 2 (early S), 4-6 (mid/late S), 8 (G2M) or 17-20 hours (G1) (Figure 2D, Figure S3B). The majority of the cells in G1 (85.3 ± 16.7%) displayed detectable *NEAT1_1* signal outside of PS, whereas in the early S, mid/late S and G2M phases *NEAT1_1* signal was found in only a very small fraction of cells (2.4 ± 3.87%; 1.63 ± 2.6% and 5.8 ±1 3.1%, respectively; Figure 2E-F and Figure S2A). Only 2.2 ± 1.4% of the cells did not display any detectable *NEAT1* staining (data not shown). We next characterized the numbers of *NEAT1* and *NEAT1_2* signals per cell in G1, S, and G2 phases. In G1 cells, the number of *NEAT1_1* foci per cell varied greatly, with an average of 24.7± 22 (Figure S2D). In contrast, the number of *NEAT1_2* detectable signals/PS fluctuated between 6 and 8.5 (Figure S2B-D).

RT-qPCR analysis showed elevated levels of total *NEAT1* in G1 compared to the other phases of the cell cycle. Since *NEAT1_2* levels remained relatively constant throughout, this is a consequence of higher *NEAT1_1* levels in this particular phase (Figure S3E-G). Accordingly, the ratio of the levels of *NEAT1_2* over *NEAT1* (*NEAT1_1* + *NEAT1_2*) (Figure S3H) in early S, mid/late S and G2 phases revolved around 1 (mean = ∼1.4 in early S and ∼0.98 in both mid/late S and G2). In contrast, this ratio was consistently smaller than 1 (mean = ∼0.18) in G1 cells, indicating that *NEAT1_1* contributes to the total levels of *NEAT1* in these cells. Moreover, in G1/G0 cells, no linear relationship could be established between *NEAT1_2* and *NEAT1* (R^2^ = 0.1141, p-value 0.259), whereas a significant positive correlation (R^2^ = 0.6488, p<0.001) was observed in S- and G2-cells. The b0 and b1 values of the equation for the linear regression (*NEAT1* = b0 + b1 * *NEAT1_2*) were nearly 0 and 1, respectively (b0 = 0.09, b1 = 1.3). These results strongly indicate that *NEAT1_1* levels drop as cells engage a new round of cell division. Similar results were obtained with another cancer cell line (HeLa cells; Figure 2E-H). We therefore concluded that *NEAT1_1* levels fluctuate during the cell cycle, whereas *NEAT1_2* levels remain relatively constant. Because *NEAT1_2* is the product of a transcription read-through event, the downregulation of *NEAT1_1* as cells engage into DNA replication cannot be due to a decrease in transcription, but must instead occur through active degradation of the transcript.

### *NEAT1_1* is degraded by the RNA exosome

In order to identify factors that contribute to the degradation of *NEAT1_1*, we mined publically available datasets and observed that *NEAT1_1*, but not *NEAT1_2*, levels were upregulated upon depletion of the exosome component RRP40 (Figure S4A). We confirmed these results by RNA-FISH (Figure 3A-B) and RT-qPCR analysis. In RRP40 KD cells, we observed a lower ratio of *NEAT1_2/NEAT1* RNA-FISH signal per nucleus, and a specific increase of the total levels of *NEAT1*, but not *NEAT1_2*, indicating that *NEAT1_1* is specifically targeted by the RNA exosome machinery (Figure 3C). Depletion of RRP40 results in the stabilization of a series of nuclear polyadenylated RNAs, which accumulate in distinct polyA+ foci (Silla et al. 2018). Combining RNA-FISH probes targeting *NEAT1* and polyA+ RNA, we detected an accumulation of *NEAT1* in the poly A+-rich foci (Figure 3D-E, S4B-C). We also noted that a pool of *NEAT1* did not overlap with poly A+ RNA, which likely represents *NEAT1_2* RNAs as these are not polyadenylated and rather stable throughout the cell cycle. To confirm this, we performed *NEAT1*- and polyA+-FISH combined with immunofluorescence analysis of the canonical PS protein NONO. Consistent with our prediction and the absence of co-staining with another PS marker, SFPQ (Silla et al. 2018), PS and *NEAT1*/polyA+ foci did not overlap (Figure 3F).

**Figure 3:**
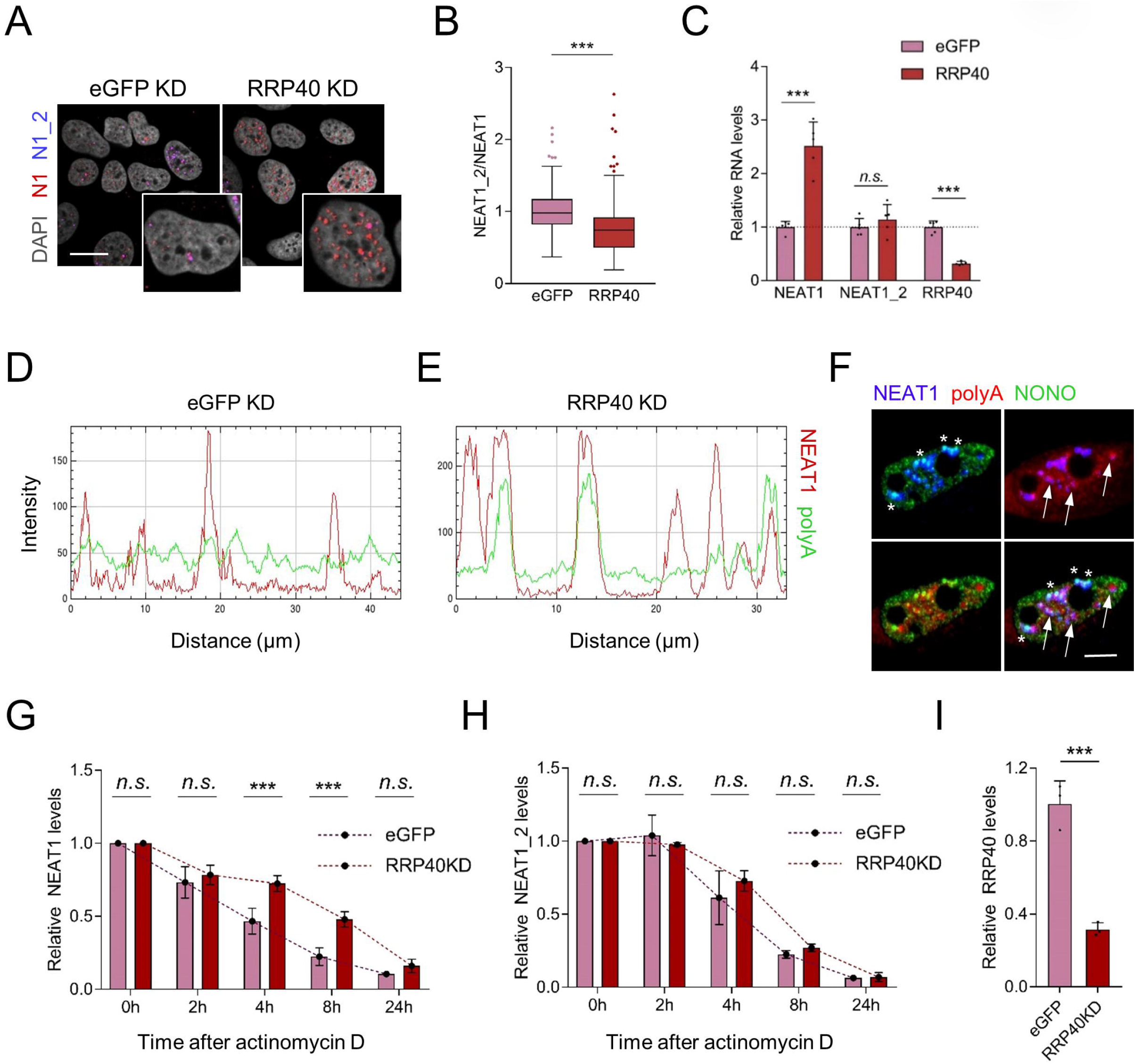
*NEAT1_1* is selectively degraded by the RNA exosome. A. Representative *NEAT1/NEAT1_2* RNA-FISH images of U2OS cells in which RRP40, a subunit of the RNA exosome complex, was knocked down. B. Tukey plots of quantified RNA-FISH signal shown as *NEAT1_2* nuclear intensity over total *NEAT1* nuclear intensity in N = 5 independent experiments. Significance was calculated by an unpaired Mann-Whitney U non-parametric test with each cell as a data point combining data from 5 independent experiments. ***, two-tailed p<0.001. C. Mean and standard deviation of relative RNA levels by RT-qPCR of the same experiments as in B (*N* = 5) in eGFP and RRP40 KD cells suggesting specific upregulation of the *NEAT1_1* isoform. Dots are individual data points. D-E. Line plots of the intensity in eGFP KD (D) and RRP40KD (E) conditions showing *NEAT1* localization in polyA+ accumulations upon exosome inhibition. F. *NEAT1* (blue) / polyA (red) RNA-FISH and IF in HeLa cells showing *NEAT1*/polyA+ accumulations in RRP40 KD conditions (arrows) are distinct from paraspeckles (*NEAT1* + NONO, asterix).G-I. Relative RNA-levels of both *NEAT1* isoforms (G) and the long *NEAT1_2* isoform specifically (H) upon addition of 2 mg/ml of the transcription inhibitor Actinomycin D 48h after eGFP or RRP40 KD in HeLa cells. I. Bar graph showing relative RNA levels of RRP40 for the experiments in G-H at time = 0h. Statistical significance was tested using a 2-way ANOVA with Dunnett’s correction for multiple testing for *N* = 3 independent experiments. Bars and error are mean and standard deviation. Individual data points are independent experiments.

To confirm that the observed upregulation of *NEAT1_1* was due to a decrease in RNA degradation rather than a transcriptional effect, we measured *NEAT1* and *NEAT1_2* levels by RT-qPCR in RRP40 depleted cells at different time points following exposure to the transcriptional inhibitor Actinomycin D. The pace at which *NEAT1_1* transcripts, but not *NEAT1_2*, decay was significantly slower in RRP40 KD cells as compared to control cells (Figure 3H-I). We concluded that *NEAT1_1* is specifically degraded by the RNA exosome.

### NEAT1_1 does not contribute to cell growth

To investigate whether *NEAT1_1* might play a role as a regulator of cell cycle progression and/or survival of G1 cells, we used CRISPR editing to delete a small regulatory region (∼140 bp) at the 3’ end of *NEAT1_1* spanning the CFIm and hnRNP K binding sites as well as the polyadenylation signal (PAS). This approach is expected to selectively delete *NEAT1_1* by allowing transcription read-through and constitutive *NEAT1_2* expression. We introduced the deletion into U2OS and two other cancer cell lines, HCT116 p53 WT and its isogenic p53 KO line Figure 4A, S5A-B), and subsequently isolated single-cell clones. PCR-based genotyping confirmed successful bi-allelic targeting of the *NEAT1_1* regulatory region, resulting in 2 wild type (WT) and four *NEAT1_1* knock out (KO) U2OS clones (Figure 4B) as well as 4 WT and 4 KO clones for each of the HCT116 cell lines (Figure S5A-B).

**Figure 4:**
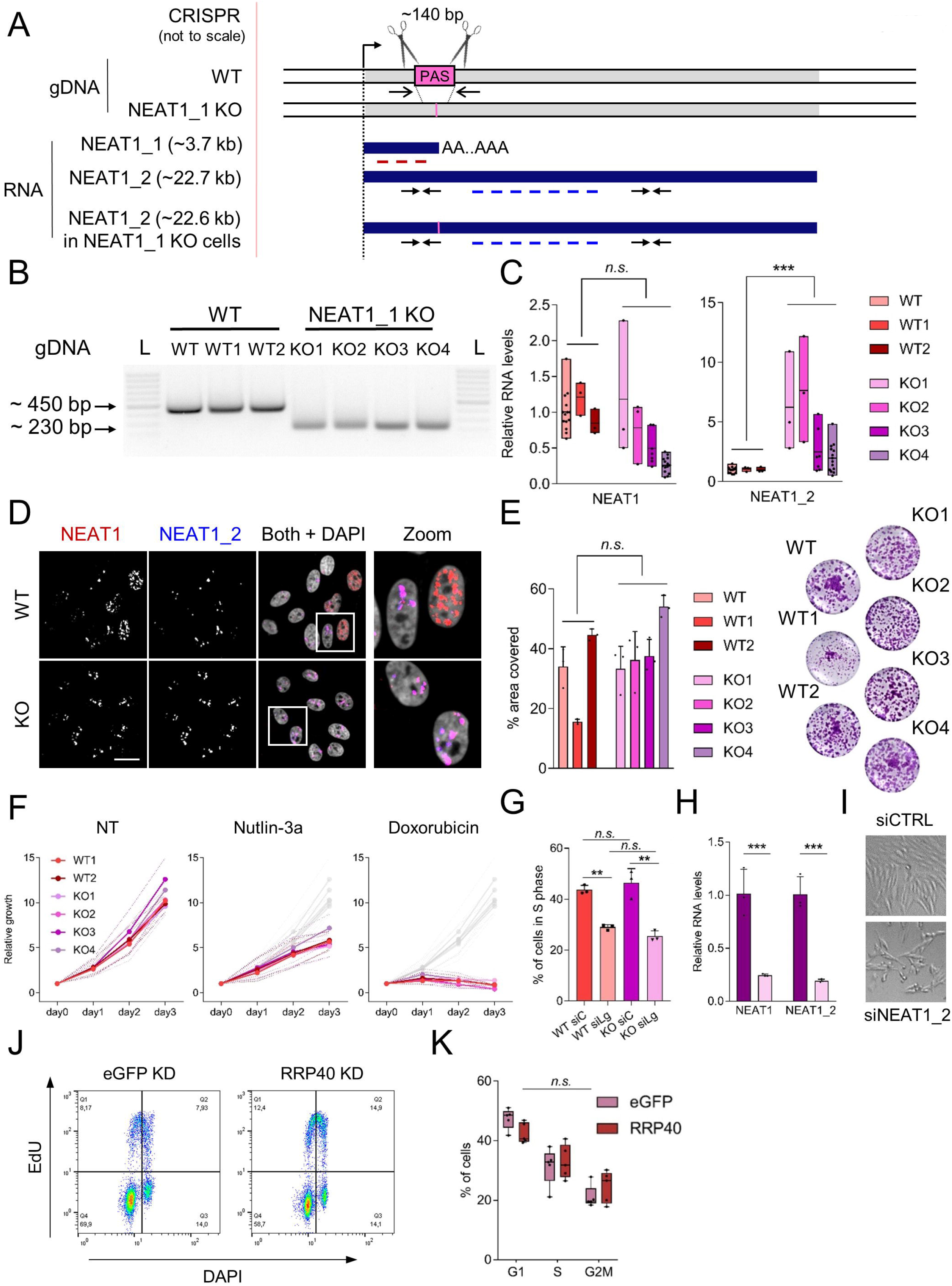
Loss of *NEAT1_1* does not impact on cell growth. A. Scheme of the CRISPR strategy used to knock out *NEAT1_1* by deletion of the regulatory sequences and polyadenylation signal at the 3’ end of the *NEAT1_1* short isoform genomic sequence and the resulting RNAs in wild type (WT) and *NEAT1_1* knock out (KO) cells. Closed arrows and dotted lines represent RT-qPCR primers and RNA-FISH probes respectively. Open arrows on the DNA sequence represent approximate locations of genotyping primers. B. Representative inverted gel image of the PCR product from U2OS gDNA used for genotyping individually isolated single cell clones after CRISPR with the primers depicted in A. L = DNA ladder. C. Relative RNA levels of *NEAT1* (total) and *NEAT1_2* specifically showing upregulation of *NEAT1_2* whereas the total levels of *NEAT1* remain the same in WT vs KO clones. The WT without a number is the mother population from which the WT and KO clones were derived. Significance was calculated using a 2-way ANOVA comparing RNA levels in WT and KO clones with Dunnett’s correction for multiple testing. ***, p<0.001 for N = at least 3 independent experiments. n.s., not significant. Dots represent data points from each independent experiment. D. Representative *NEAT1/NEAT1_2* RNA-FISH image of WT and *NEAT1_1* KO cells showing the complete loss of *NEAT1_1* upon polyA site knockout. Scale bar, 10 µm. E. Quantification of % area covered (left) and representative images (right) of colony assays 14 days after seeding 2000 cells per well in *N* = 3 independent replicates, 3 wells per replicate each. Statistical testing was done using a 1-way ANOVA. n.s.: not significant. Dots represent average of 3 wells of the independent experiments. Bars are mean + standard deviation. F. Short term growth measured by WST-1 relative to day 0 (1 day after seeding) in non-treated (NT), 10 µM Nutlin-3a and a low dose of doxorubicin (150 ng/ml) in *NEAT1_1* WT and KO clones. All data is the average of *N* = 3 independent experiments. Standard error is depicted as dotted lines above and below the data points. In Nutlin-3a and doxorubicin conditions, values in non-treated conditions are shown as light grey lines in the back of the graph. All data is not significant as tested by 2-way ANOVA with Dunnett’s correction for multiple testing in the different time points. G. Quantification of % of EdU positive cells (S phase) in flow cytometry upon CTRL (siC) and *NEAT1_2* (siLg) knockdown in WT and KO clones. N = 3 independent experiments. Significance was determined using 2-way ANOVA with Dunnett’s correction for multiple comparisons. ***, p<0.001. n.s.: not significant. I. representative image of siCTRL and si*NEAT1_2* KO cells showing decreased cell density 48h after transfection. J. Representative flow cytometry graphs of EdU/DAPI staining in eGFP and RRP40 KD conditions. K. % of cells in G1, S and G2M phases of the cell cycle upon eGFP and RRP40 KD as in Figure 3A-C. *N* = 5 independent experiments. Non-significance was determined using a 2-way ANOVA comparing eGFP and RRP40 conditions in each of the phases with Dunnett’s correction for multiple testing.

To establish that the engineered cells did not express *NEAT1_1*, we quantified total *NEAT1* and *NEAT1_2* levels using RT-qPCR. Whereas relative *NEAT1* levels did not change in the PAS KO clones, *NEAT1_2* levels were increased, consistent with the prediction that all initiated transcripts contribute to expression of the long isoform (Figure 4C, S5E-F). In agreement, RNA-FISH analysis did not detect *NEAT1_1* in the nucleoplasm of the KO clones (Figure 4D, Figure S6C-D). Notably, PS integrity was preserved in the KO cells, as evidenced by their co-staining with the PS marker NONO and the *NEAT1* RNA FISH probe sets (Figure S6).

We next assessed whether the *NEAT1_1* deletion affects cell growth and proliferation. Long-term growth assays indicated that *NEAT1_1* KO cells proliferated at a similar rate as the WT controls (Figure 4E, Figure S5J-M), which was confirmed in a short-term growth assay (WST-1) and following exposure to Nutlin-3a or a low dose of the DNA damaging agent doxorubicin (Figure 4F, Figure S5G-I). This is in contrast to specific transient depletion of *NEAT1_2*, which sensitized cells to these agents (Adriaens et al. 2016). These results thus indicated that *NEAT1_1* is not required for the two-dimensional growth and proliferation of cancer cell lines.

To further assess *NEAT1_1* and *NEAT1_2* independent functions, we knocked down *NEAT1_2* in the PAS KO cells and analyzed their cell cycle distribution and growth properties. *NEAT1_2* knockdown induced a similar decrease in EdU-positive cells as it did in WT, cycling cells (Figure 4G-H). Cell density was also markedly decreased upon *NEAT1_2* KD (Figure 4I). These data indicated that *NEAT1_1* does not contribute to the ability of *NEAT1_2* to preserve the genomic integrity of cancer cell lines.

Moreover, we could not identify cell cycle defects in RRP40-depleted cells, indicating that *NEAT1_1* accumulation in polyA+ foci does not affect cell division (Figure 4J-K).

### NEAT1_1 depletion does not overtly impact on the cellular transcriptome

It has been proposed that *NEAT1* regulates cellular gene expression by localizing to the transcription start sites of actively transcribed genes (West et al. 2014). In order test whether *NEAT1_1*, which is found prominently in the nucleoplasm of G0 and G1/S cells (Figure 5A), modulates transcription we profiled the transcriptome of PAS KO and WT ctrl cells by RNAseq. We detected on average 18.030 (G0) and 17.250 (G1S) expressed genes (figure 5B), of which only 156 (∼0.86%) and 23 (∼0.13%), respectively, were significantly differentially expressed (DE) in the PAS KO compared to WT cells (Figure 5C-F). Gene ontology analysis did not identify particular pathways or biological processes affected by the depletion of *NEAT1_1*. Thus, although *NEAT1_1* is highly expressed in G0/G1 cells, its loss does not significantly impact the overall gene expression profiles of these cells.

**Figure 5:**
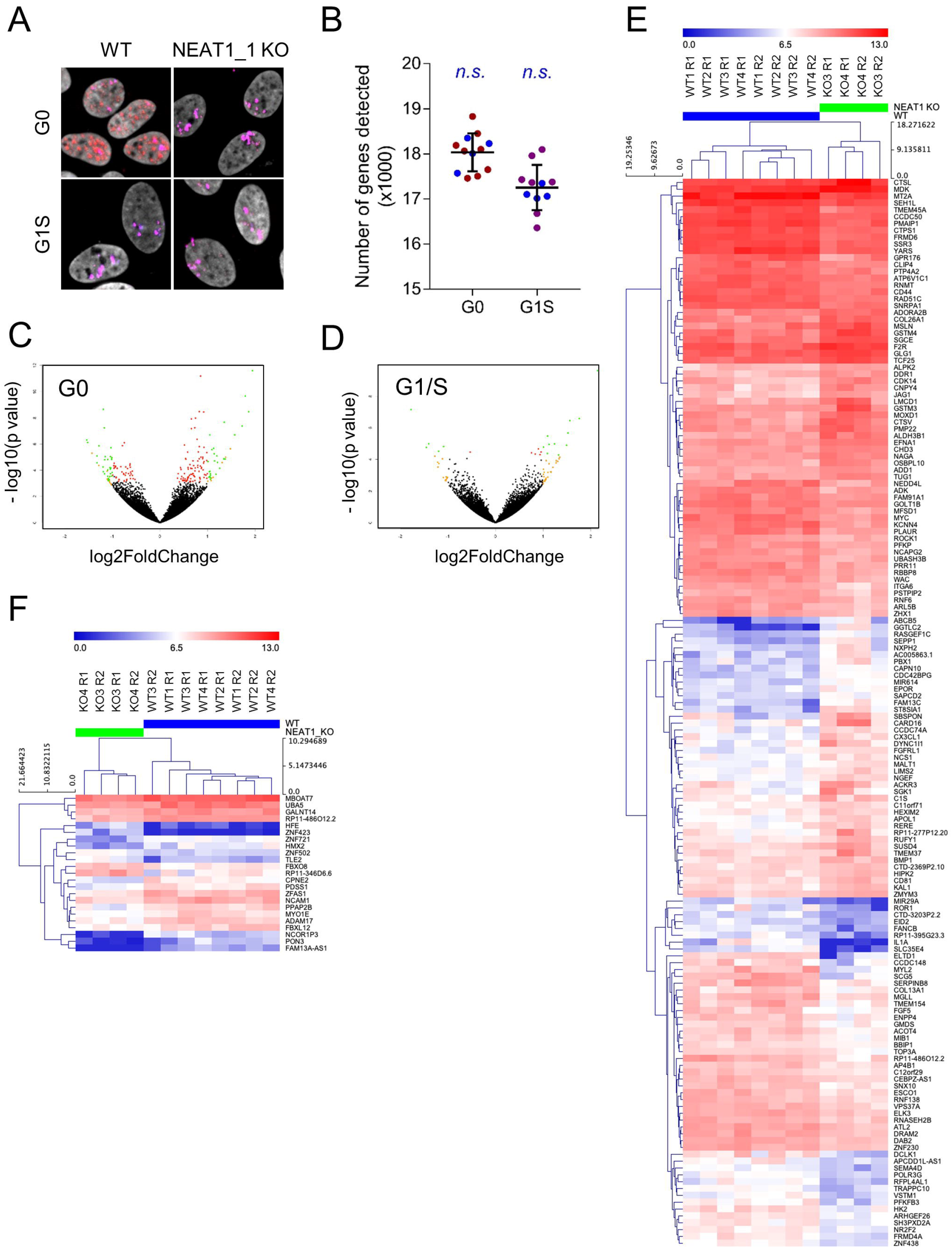
*NEAT1_1* KO only causes limited changes in gene expression. A. Representative *NEAT1/NEAT1_2* FISH in G0 and G1S conditions as in Figure 2 A-C in WT and *NEAT1_1* KO cells used for the Smartseq2 RNA sequencing experiment. B. Number of genes detected in G0 and G1s conditions. Red/pink dots are WT conditions, blue dots represent KO conditions. Significance was tested using a two-sided unpaired t-test comparing the number of genes detected in WT and KO conditions. C-D. Volcano plot of gene expression changes (-log2) in G0 (C) and G1S (D) plotted against their p-value (−log10). Dots are color coded red if the adjusted p<0.05, orange if log2 fold change >1 and green if both. E. Hierarchical clustering of G0 samples based on 156 unique differentially expressed genes (FC>1.5, p-adj<0.05). F. Hierarchical clustering of G1/S samples based on 23 unique differentially expressed genes (FC>1.5, p-adj.<0.05).

### Neat1_1 KO mice do not exhibit lactation nor fertility defects

To further explore a physiological function of *Neat1_1* in normal cells and in the relevant *in vivo* context, we generated a *Neat1_1*-specific KO mouse strain using a strategy similar to the one used to knock out *NEAT1_1* in cells. In brief, 39 base pairs surrounding the PAS of *Neat1_1* were excised using CRISPR/Cas9 in mouse Embryonic Stem Cells (mESCs), and a mouse strain deficient for *Neat1_1* generated (S. Nakagawa and T. Hirose, manuscript in preparation). PAS KO mice were born at the expected Mendelian ratios (Figure 6D) and did not exhibit the lactation defect previously observed in *Neat1* full KO mice (Standaert et al. 2014). We weighed pups born in nests from WT, *Neat1* full KO, *Neat1* heterozygous, PAS heterozygous and PAS KO mothers at three and six weeks of age and confirmed that full KO females were unable to successfully nurture their pups. In contrast, PAS KO females gave birth to normally sized nests and all offspring developed and gained weight normally (Figure 6E-F).

**Figure 6:**
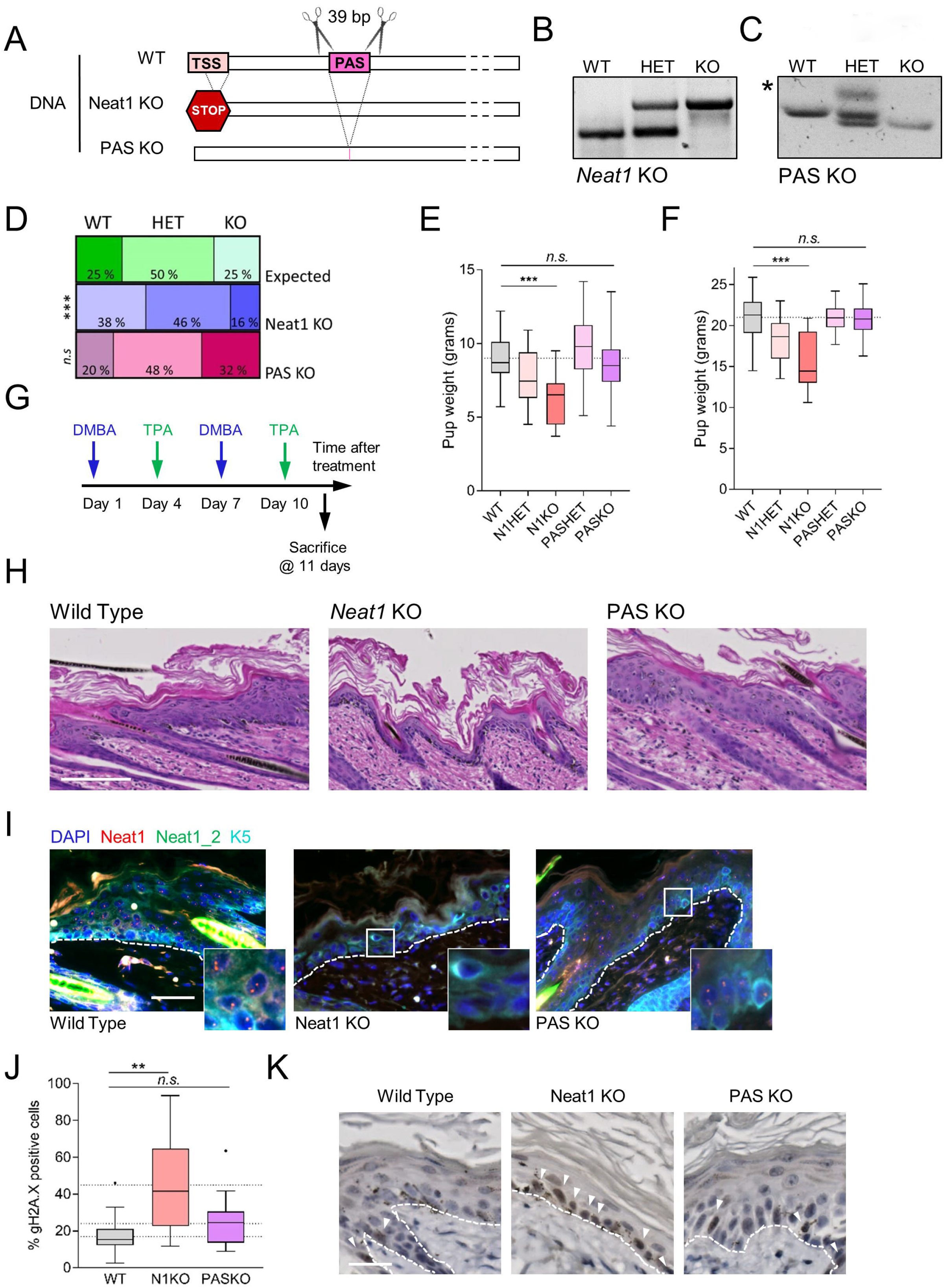
Comparison of *Neat1* versus *Neat1_1* KO phenotypes. A. CRISPR strategy to knock out *Neat1_1* in mouse embryonic stem cells resulting in a 39 base pair deletion spanning the *Neat1_1* polyadenylation signal (PAS) and strategy to knock out both isoforms as described in Nakagawa et al., 2011. B. Genotyping of *Neat1* full KO mice. C. Genotyping of *Neat1_1* (PAS) KO mice. The asterix indicates an unspecific band in the heterozygous sample. D. Genotype distribution of pups born from heterozygous parents in full *Neat1*KO (middle bar) and PAS KO nests (lower bar) as compared to the expected mendelian ratios (upper bar). *N* = 53 and 15 litters for *Neat1* and PAS KO, respectively. P-values were calculated using the chi-squared test. ***, p<0.001. n.s., not significant E-F. Pup weight of offspring from females with the respective genotypes at 3 (E) and 6 (F) weeks of age. Tukey plots of pups from *N* = between 4 and 19 females per genotype. Statistical significance was calculated using 2-way ANOVA with Dunnett’s correction for multiple testing. ***, p<0.001. n.s., not significant. G. Strategy and time line for short-term DMBA/TPA carcinogenesis protocol. H. Representative H&E staining of back skin sections from mice treated as in G. Scale bar, 50 µm. I. *Neat1* RNA-FISH and Keratin 5 immunofluorescence on back skin section of mice treated with DMBA and TPA as in G. J-K. Quantification (J) and representative images (K) of sections from DMBA/TPA treated back skin stained with the DNA damage marker γ-H2A.X of *N* = at least 3 mice per genotype according to the method described in Adriaens et al., 2016. The Tukey plot in J graphs individually quantified pictures. Statistical testing was done on biological replicates (averages for individual mice) using a 1-way ANOVA with Sidak’s correction for multiple testing. **, p<0.01. n.s., not significant. White arrows in K indicate γ-H2A.X positive cells, white dotted lines separates the dermis from the epidermis. Scale bar, 20 µm.

To further study the impact of the mutation on the growth of normal cells *in vitro*, we produced PAS KO MEFs. Despite an increase in *Neat1_2* levels (Figure S7A-C), passage 3 MEFs derived from the PAS KO mice grew similarly to WT MEFs. Full *Neat1* KO also grew similarly to WT fibroblasts (Figure S7D-E) (Nakagawa et al. 2011; Adriaens et al. 2016).

### Mouse Neat1_1 does not contribute to DNA damage induction and reduced growth during skin carcinogenesis

The skin of *Neat1* KO mice exhibits an exacerbated sensitivity to DNA damage and, thereby, an increased resistance to DMBA/TPA-induced skin hyperplasia and tumorigenesis (Adriaens et al. 2016). To test whether *Neat1_1* mice exhibit a similar phenotype, we subjected these mice to the DMBA-TPA protocol and assessed PS formation, measured hyperplasia and accumulation of DNA damage in their treated back skin. We found that in our short-term protocol (11 days of treatment), both PAS KO and WT cells displayed moderate to severe hyperplasia (Figure 6H) and abundant PS (Figure 6I). In contrast, back skin of *Neat1* KO mice neither displayed PS nor marked hyperplasia (Figure 6H-I). In addition, it showed a significant increase in persisting DNA damage in the treated regions as compared to the skin of WT and PAS KO animals (figure 6J-K). We concluded that the phenotypes observed in *Neat1* KO mice are attributable to the loss of the long *Neat1_2* isoform and, thereby likely to be a consequence of loss of PS nuclear bodies.

## Discussion

In this work, we demonstrate that the expression levels of the two *NEAT1* isoforms are dynamically regulated during the cell cycle. We observed that the short isoform, *NEAT1_1* is highly expressed in G0/G1-phase of the cell cycle and that, in line with previous findings (Li et al. 2017), it localizes prominently outside of PS. We also observed that *NEAT1_1* levels drop abruptly as cells transit from the G1 to the S phase. This is consistent with the observation that *NEAT1_1* is detected at high levels in terminally differentiated cells in most tissues (Nakagawa et al. 2011). In contrast, *NEAT1_2* levels remain relatively constant throughout the cell cycle and consequently *NEAT1_2* is the only detected *NEAT1* isoform in S-phase and onwards in these cells, which is in keeping with previous data reporting the presence of PS in amitotic (interphase) cells as evidenced by the typical clustering of canonical PS protein p54nrb/NONO in their nuclei (Fox et al. 2005).

It had been proposed that the short *NEAT1* isoform is recruited into PS to support their stability and/or functions (Souquere et al. 2010; Mao et al. 2011; West et al. 2016). In disagreement with this possibility, we show that when both isoforms are co-expressed in G0/G1, a large fraction of the *NEAT1_1* isoform transcripts localize outside of PS. In cells residing in other phases of the cell cycle, *NEAT1_2* is the only isoform present and therefore PS are, by and large, *NEAT1_1*-free in these cells. Our data is, however, in line with previous quantifications of *NEAT1_1* RNA levels, indicating that, on the basis of stoichiometry, *NEAT1_1* transcripts are not likely to locate to PS, or at least not in significant amounts (Chujo et al. 2017; Li et al. 2017). Our data also suggest that PS can be assembled in the absence of *NEAT1_1*. This observation resonates with previous work showing that *NEAT1_2* is the *NEAT1* isoform required for PS assembly (Sunwoo et al. 2009) and that *NEAT1_1* expression alone is not sufficient to rescue PS formation upon *NEAT1_2* ablation (Sasaki et al. 2009; Mao et al. 2011).

Our data demonstrate that the *NEAT1_1* transcript is actively degraded as cells commit to divide. We provide evidence that this process is mediated by the main RNA degradation machinery in the nucleus, namely the RNA exosome (Vanacova and Stef 2007). Knockdown of one of its core components, RRP40, led to the specific accumulation of the *NEAT1_1* short isoform within nuclear bodies containing persistent polyA+ RNA transcripts (Silla et al. 2018). We provide evidence that the RNA exosome mainly targets *NEAT1_1* (and not *NEAT1_2*). Moreover, *NEAT1_1*/polyA+-containing foci did not overlap with PS, consistent with *NEAT1_1* being spatially distinct.

Our observations that the evolutionarily conserved *NEAT1_1* isoform is (i) produced at high levels in most-if not all- resting cells (as well as in cancer cells in G1), and (ii) gets actively degraded as cells commit to divide, suggested a putative role for *NEAT1_1* in the regulation of the cell cycle. However, silencing of RRP40 did not overtly perturb progression of the cell cycle. Consistently, previous reports have shown that modulating levels of neither RRP40 and its targets (Graham et al. 2009; Zinder and Lima 2017) nor *NEAT1_1* affect cell cycle progression (reviewed in Yu et al. 2017). More strikingly, *NEAT1_1* isoform specific KO cells grew and responded to stress comparably to WT control cells. Moreover, PS formed normally in these cells and the phenotypes observed upon silencing of *NEAT1_2* in various cancer cell lines were also observed in *NEAT1_1* KO cells when co-depleted for *NEAT1_2*. Together, these data indicated that *NEAT1_2* functions independently of *NEAT1_1* and that *NEAT1_2* can promote PS formation in absence of the short *NEAT1* isoform.

Our data demonstrating that *NEAT1_1* localizes outside of PS is consistent with another report showing that there is a fraction of *NEAT1* that localizes diffusely throughout the nucleoplasm (Li et al. 2017) and the possibility that *NEAT1* binds active transcription start sites in euchromatin (West et al. 2014). However, only minor changes in gene expression patterns were observed in *NEAT1_1 KO* cells, indicating that *NEAT1_1* does not directly impact on transcription. Instead, we hypothesize that its enrichment in euchromatin might have spatial, physical grounds (i.e. diffusion is easier in euchromatin due to a decrease in molecular crowding) rather than a specific functional role in modulating gene expression. Moreover, in contrast to *Neat1* KO mice (Nakagawa et al. 2014), their *Neat1_1* counterparts were born at the normal Mendelian ratios. Pups born from *Neat1_1* KO females were not significantly smaller than those from wild type and heterozygous mothers. This is in contrast to our previously published observation that *Neat1* KO females cannot nurture their pups properly owing to mammary gland formation and lactation defects (Standaert et al. 2014). Similarly, *Neat1_1* KO mice did not exhibit any of the phenotypes observed in *Neat1* KO mice exposed to a 2-step skin carcinogenesis protocol (Adriaens et al. 2016).

Taken together, our data suggest that *NEAT1_1* is a non-functional, non-essential isoform in both resting and proliferative cells, at least in the interrogated experimental conditions. Is *NEAT1_1* then just a non-functional byproduct of the *NEAT1* locus? Perhaps active transcription of the *NEAT1* locus ensures that cells can rapidly switch to *NEAT1_2* production in response to stress and thereby the formation of PS. Constant synthesis of a nonfunctional *NEAT1_1* transcript would therefore be the price that cells have to pay to be able to quickly engage a PS-dependent survival pathway when exposed to deleterious stimuli. However, It cannot be excluded that *NEAT1_1* does exert a function in very specific stress and/or pathophysiological conditions, for instance during viral infections (Saha et al. 2006; Zhang et al. 2013; Imamura et al. 2014; Ma et al. 2017; Morchikh et al. 2017; Wang et al. 2017). The enclosed described *Neat1_1* mouse model will be a valuable tool to test this possibility.

## Materials and methods

### Cell lines, culture methods and cell synchronization

U2OS, HeLa and HCT116 WT and p53KO isogenic cell lines were obtained from the LGC ATCC and maintained in DMEM (ThermoFisher Scientific Cat. No. 21885025) plus 10% fetal bovine serum (FischerScientific Cat. No. 10270106). None of the cell lines used were reported in the ICLAC database of commonly misidentified cell lines. All cell lines were tested monthly for mycoplasma contamination and found negative. After their initial purchase, cell lines were not further authenticated. For synchronization in G0, the cells were washed with PBS 24 hours after plating and media were replaced with media containing no serum for three days. For G1/S synchronization, G0 cells were released in 20% serum plus 5 µg/ml Aphidicolin (Sigma, A0781) for 24h. For G1, S and G2 synchronization, media was replaced with media containing 2 mM thymidine (Sigma, T1895) for 12h, released in normal growth media for 12h and then grown again in thymidine-containing media for 12h prior to release in normal media and harvesting at the indicated time points. For knockdown experiments with siPOOLs (siTOOLS Biotech), 25 nM siRNA was transfected using Lipofectamine RNAiMAX (ThermoFisher Scientific, Cat. No. 13778075) as previously described (Adriaens et al. 2016).

### Generation and culture of Mouse Embryonic Fibrobasts

Mouse Embryonic Fibroblasts were generated from plug-checked pregnant females at E12.5. The embryos were removed from the uterus and internal organs were discarded with sterile forceps. Heads were used for genotyping. The remainder tissue was pipetted up and down in sterile PBS several times to obtain single cell suspensions before transfer to tissue culture dishes with DMEM containing 10% serum, 1% penicillin/streptomycin (Invitrogen, Cat. No. 15140122) and 50 µM β-Mercaptoethanol (ThermoFisher Scientific, Cat. No. 31350010). The cells were passaged twice before all experiments were performed at passage 3.

### RNA-FISH

Dual RNA-FISH was performed according to the Stellaris RNA-FISH protocol essentially as described in (Adriaens et al. 2016) but with probesets against both (Cat. No. VSMF-2246-5 and VSMF-2247-5) or *NEAT1_2* specifically (Cat. No. VSMF-2251-5) in human cells and against *Neat1* both (Cat. No. VSMF-3030-5) or *Neat1_2* specifically (Cat. No. VSMF-3035-5) in mouse cells and mouse tissues. Cells were generally plated on 11 mm coverslips in 6-well plates to allow for concomitant RT-qPCR analysis, cell cycle analysis and RNA-FISH. Costaining with the paraspeckle marker p54nrb/NONO (Souquere et al. 2010) was done after the ethanol permeabilization step of the RNA-FISH protocol. Briefly, the cells were washed once with PBS followed by a 5’ permeabilization step with 0.5% Triton-X100 in PBS. Then, the cells were incubated with a 1/1000 dilution of the antibody in DAKO antibody dilution reagent for 1h at room temperature followed by three washes in 0.05% Tween-20 in PBS and staining with the secondary antibody (Life Technologies, anti-mouse A488) in DAKO for 1 hour at RT. After two short washes in 0.05% Tween-20 in PBS, the cells were incubated with Wash Buffer (2x SSC, 10% v/v Formamide (Sigma Aldrich, Cat. No. F9037) and the RNA-FISH protocol was continued as described. Hybridization buffer was made using the same formula as wash buffer, adding 10% w/v Dextran (Sigma Aldrich, Cat. No. D8606) and probes at a final concentration of 25 nM. Images were acquired with a Nikon A1 confocal microscope acquired through a Hercules grant type 1 AKUL/09/037 and processed for overlay and brightness and contrast adjustments using ImageJ. RNA-FISH images from mouse back skin tissue were acquired with a ZEISS Axio Scan Z1 microscope using 20x and 40x objectives followed by stitching of the continuous fields using ZEN2 software.

### Image analysis

Confocal images were quantified using FIJI software (ImageJ 1.51p. Java version 1.8.0_66, 64-bit, National Institutes of Health, USA). To determine the number of cells that display *NEAT1_1* outside of paraspeckle nuclear bodies, we processed the raw images with the Speckle Inspector plugin on each channel after thresholding, with a minimal speckle size 2 pixels, within the nuclear region delineated by the DAPI channel. The number of spots in the *NEAT1_2* channel was subtracted from the number of spots per cell in the *NEAT1* channel. If the outcome of the subtraction was larger than 3 (arbitrary error margin: 0 (expected) +3 to account for accidental background spots in the Q570 channel), we considered that the cells contained detectable *NEAT1_1* outside of the paraspeckles. The total numbers of cells were determined using the Cell Counter plugin. Nuclear RNA-FISH intensity (Figure 3G, 4B) was calculated by thresholding, filling holes and watershed of the DAPI channel, and determination of the nuclear intensity in the *NEAT1* and *NEAT1_2* channels per cell via the “send to” functionality in Set Measurements before Counting Particles. Tthe percentage of cells containing detectable *NEAT1_1* is represented on the left y-axis, whereas the percentage of cells that only displayed paraspeckles was represented on the right y-axis.

To find the ratios per cell of *NEAT1_2* signal over total *NEAT1* signal, we used the FIJI function “Set Measurements” as above to redirect DAPI-thresholded images to the respective *NEAT1* (Quasar 570, measured in the red channel and represented in red) and *NEAT1_2* (Quasar 670, measured in the far red channel and represented in blue) channels to obtain their relative intensities, which were then plotted per cell as *NEAT1_2/NEAT1* relative integrated density per cell.

### RNA isolation and RT-qPCR

RNA isolation, generation of cDNA and RT-qPCR were performed essentially as described in (Adriaens et al. 2016). Briefly, after lysis, the cell-lysis buffer mixture was heated for 10 minutes at 55°C according to the protocol described in Chujo et al., 2017. Then, total RNA was isolated using the NucleoSpin RNA kit (Macherey Nagel, Cat. No. 740955), including rDNAse treatment for 15 minutes according to the manufacturer’s instructions. The RNA was reverse transcribed using the ThermoFisher Scientific High Capacity cDNA Reverse Transcription Kit (Cat. No. 4368813). RT-qPCR was performed with GC Biotech SensiFast SYBR No-Rox (Cat. No. BIO-98020) and run on a Roche LightCycler-480-384. For normalization, the geometric mean of the two most stable reference genes out of at least three was calculated using geNorm in qBase+ Software (Biogazelle, Zwijnaarde, Belgium - www.qbaseplus.com). RT-qPCR primer sequences were as follows: *NEAT1* fw: 5’-GGAGAGGGTTGGTTAGAGAT-3’; *NEAT1* rev: 5’-CCTTCAACCTGCATTTCCTA-3’; *NEAT1_2* fw: 5’-GGCCAGAGCTTTGTTGCTTC-3’; *NEAT1_2* rev: 5’- GGTGCGGGCACTTACTTACT-3’; *CDKN1A* fw: 5’-AGCAGAGGAAGACCATGTGGA-3’; *CDKN1A* rev: 5’- AATCTGTCATGCTGGTCTGCC-3’; *UBC* fw: 5’- ATTTGGGTCGCGGTTCTTG-3’; *UBC* rev: 5’-TGCCTTGACATTCTCGATGGT-3’; *TBP* fw: 5’- CGGCTGTTTAACTTCGCTTC-3’; *TBP* rev: 5’- CACACGCCAAGAAACAGTGA-3’; *B2M* fw: 5’-TGCTGTCTCCATGTTTGATGTATCT-3’; *B2M* rev: 5’-TCTCTGCTCCCCACCTCTAAGT-3’; *HPRT1* fw: 5’-TGACACTGGCAAAACAATGCA-3’; *HPRT1* rev: 5’- GGTCCTTTTCACCAGCAAGCT-3’; *mNeat1_2* fw 5’-GCTCTGGGACCTTCGTGACTCT-3’; *mNeat1_2* rev 5’-CTGCCTTGGCTTGGAAATGTAA-3’; *mNeat1* fw 5’- TTGGGACAGTGGACGTGTGG-3’; *mNeat1* rev 5’-TCAAGTGCCAGCAGACAGCA-3’; *mHmbs* fw 5’-GCGGAGTCATGTCCGGTAA-3’; *mHmbs* rev 5’- GTGGTGGACATAGCAATGATTT-3’; *mGapdh* fw 5’-AGGTTGTCTCCTGCGACTTCA-3’; *mGapdh* rev 5’-GGTGGTCCAGGGTTTCTTACTC-3’. For correlation analysis, primer efficiencies were calculated in qBase+ by combining cDNA of each of the tested samples and producing a serial dilution (1, 0.5, 0.25, 0.125, 0.0625, 0.03125) to be run simultaneously with the individual samples.

### Cell cycle analysis

Cell cycle analysis was performed by pulsing the cells with 10 µM of EdU for 30 minutes before harvesting and trypsinization, or via DNA profiling alone against a non-synchronized control to identify 2N and 4N populations. For EdU staining, the cells were washed in cold PBS + 10% serum to inactivate the trypsin, collected by centrifugation and fixed for 15 minutes with 4% PFA in PBS. After two washes in PBS, the cells were stored overnight in 15 mL 0.01% Triton-X100 at 4°C. To detect cells in S phase, the cells were subjected to a modified Click-IT reaction protocol (Click-iT® EdU Alexa Fluor® 488, Cat. No. C10420). Briefly, the cells were collected by centrifugation, the supernatant was discarded, and they were incubated in 50 µL Click-IT reaction cocktail (43.75 µL PBS + 1 µL CuSO4 100 mM + 5 µL 100 mM Ascorbic Acid + 0.25 µL A488 Azide dye) for 50 minutes. The cells were then washed once in PBS and resuspended in 300 µL DAPI staining buffer (5 µg/mL DAPI in PBS with 0.1% w/v BSA) before analysis on a MACSquant® VYB flow cytometer (Miltenyi Biotec, Germany).

### CRISPR/Cas9 plasmid construction

Guide RNAs were designed for the 3’ regulatory region of *NEAT1_1* using http://crispr.mit.edu. Five µg of plasmid pSpCas9(BB)-2A-GFP (pX458) (Addgene, Cat. No. 48138) was digested with BbsI/BpiI (ThermoFisher Scientific) and purified using a NucleoSpin® Gel and PCR Clean-up Kit (Machery Nagel, Cat. No. 740609) followed by In- fusion cloning of annealed gRNA oligos with 20 nucleotide overhangs on both sides (IDT) with sequences according to the manufacturer’s instructions (Takara Bio Cat. No. 121416). The gRNA sequences used to generate 4 different Cas9 targeting plasmids were: upstream Guide #1 5’-GTGTATTAGTCACGCATGTATGG-3’ quality score 89; upstream Guide #7 5’-GTACTGGTATGTTGCTCTGTATGG-3’, quality score 70; downstream Guide #1 5’-GTACATCCAAAGTCGTTATGAAGG-3’, quality score 90; downstream Guide #4 5’- GCGTTATGAAGGCAATGTGATAGG-3’, quality score 70. Following in-fusion cloning, the plasmids were transformed into competent bacteria (DH5α) grown on ampicillin plates. A colony PCR was performed to check for the correct insertion of the gRNA sequence using GoTaq Green Mastermix (Promega, Cat. No. M712) and primers 5’- GAGGGCCTATTTCCCATGATT-3’ (fw) and 5’-AAAAAAGCACCGACTCGGTGCCA-3’ (rev). Positive clones were further expanded and their inserted sequences were verified with Sanger sequencing at the VIB Genomic Service Facility, Belgium using the same primers.

### Generation of *NEAT1_1* KO cells

Once we obtained the desired Cas9/gRNA constructs, we transfected cells plated in 10 cm dishes with 10 µg of downstream and 10 µg of upstream plasmid (Combination dG1/uG1 for U2OS and HCT116 and dG4/uG7 for HCT116) using a standard transient overexpression protocol with Lipofectamine 2000 reagent according to the manufacturer’s instructions (ThermoFisher Scientific, Cat. No. 11668019). 48h after transfection, we sorted the cells for GFP expression using a S3™ Sorter (Bio-Rad Laboratories, USA) and diluted the cells at 0.5 cells/100 µL into 96 well plates. After 2 weeks of culture, we visually inspected the wells and selected those containing a single clone. These were collected and replated in duplicate. The cells in one of the two wells were then lysed and subjected to PCR analysis to determine their *NEAT1_1* genotype with primers 5’-CGTTGGGATCTTTCTGTCT-3’ (fw) and 5’- GCTCTCCTACATGGCCTTAAT-3’ (rev). These primers were also used for Sanger sequencing to characterize the repair on each allele in homozygous *NEAT1_1* KO clones. Several homozygous WT and homozygous KO clones were then selected and expanded into new cell lines from the remaining wells.

### Cell growth assays

To determine long term cell growth, cells were plated at the indicated densities in three wells per cell line per experiment and grown for 10 or 14 days. They were washed twice in cold PBS, followed by staining for 15 minutes with 0.5% Crystal Violet (Sigma Aldrich, Cat. No. C6158) in 20% Methanol/80%H20. The plates were washed and rinsed in tap water and the % area covered of the wells was quantified using FIJI. For short term growth assays, 1500 cells were plated followed by incubation with WST-1 reagent (Roche, Cat. No. 05 015 944 001) and measurement of the luminescence with a VICTOR X3 Multilabel Plate Reader (PerkinElmer) at the indicated time points. Cells were treated with 10 µM Nutlin-3a (Sigma Aldrich, Cat. No. SML05080) or 150 ng/ml doxorubicin (Sigma Aldrich, Cat. No. D1515).

### RNA sequencing

Total RNA was extracted as described above using the NucleoSpin RNA kit (Machery Nagel, Cat. No. 740955). The RNA integrity was monitored using Bioanalyzer analysis (Agilent, RIN: 9.7-10). About 500 pg of RNA per sample was reverse-transcribed and amplified using a modified SMARTseq2 protocol (Rambow et al. 2018). Prior to generating sequencing libraries using the NexteraXT kit (Illumina, Cat. No. FC-131-10), cDNA profiles were monitored using the Bioanalyzer. Sequencing was performed on a Nextseq500 platform (Illumina, SE75bp). Reads were then mapped to the human genome (hg19) using STAR (2.4.1b) and quantified with Subread (1.4.6-p2). Differential analyses between *NEAT1_1* KO and WT samples (during G0 and G1/S) were executed using the DeSeq2 pipeline. Samples were grouped using hierarchical clustering (Euclidean distance) based on differentially expressed genes (MeV4_8_1).

### KO mice

*Neat1* KO*, Neat1_1* KO and WT mice were maintained on a pure C57BL/6J background in a certified animal facility at KU Leuven Campus Gasthuisberg, Leuven, Belgium. They were maintained on a 12/12h light/dark cycle and had access to food and water *ad libitum*. All animal experiments were carried out in accordance with the guidelines of the Ethical Committee University of Leuven Animal Care and Use under project license 089/2013. Full *Neat1* KO mice were described previously (Nakagawa et al. 2011) and genotyped with primers 5’- GGTGACGCGACACAAGAGTA-3’ (fw), 5’-AAATGTGAGCGAGTAACAACCC-3’ (rev WT) and 5’- CTGTGAAACTTGTGCCCTCC-3’ (rev KO) giving rise to PCR products of 612 base pairs (*Neat1* KO) and 336 base pairs (WT). *Neat1_1* KO mice were generated by S. Nakagawa and T. Hirose using a similar CRISPR-Cas9 strategy as described for the cancer cells above generating a 39 base pair deletion of the polyadenylation signal (5’- ACAGCAAAATAAAGGTTTGAGATTGAAGCTTCTTAGAAT-3’) and genotyped with primers 5’-GCAAAGTGACAGAGGTCGAGA-3’ (fw) and 5’-AGGCAAAGTGACAGAGGTCG-3’ (rev) (WT allele: 145 base pairs; KO allele, 106 base pairs) (Unpublished, manuscript in preparation). In order to test for lactation defects, mice with mothers from the indicated genotypes were weighed at 3 and 6 weeks of age (Standaert et al. 2014). Ratios of animals born at indicated genotypes to test against expected Mendelian genotype ratios were calculated from heterozygous x heterozygous parents in both colonies.

### DMBA/TPA protocol

The DMBA/TPA protocol was performed as described in (Adriaens et al. 2016).

### H&E and immunohistochemistry

Immunohistochemistry and quantification of images was performed as described in (Adriaens et al. 2016) using antibodies against γ-H2A.X (Cell Signaling, 2577; 1/1,400) and Keratin 5 (rabbit polyclonal anti-keratin 5; Covance, PRB-160P-0100; 1/1,000). For immunofluorescence the secondary antibody was anti-Rabbit-A488 (Life Technologies). Images were acquired with a ZEISS Axio Scan Z1 microscope using 20x and 40x objectives followed by stitching of the continuous fields using ZEN2 software.

## Author contributions

CA designed the study, performed all experiments and analyzed all data apart from the RNA sequencing analysis. FR performed RNA sequencing analysis. GB performed experiments. TS provided images from RRP40 KD conditions and found the link with *NEAT1_1* specific degradation. TH, MM, TC, AH and SN provided reagents and made the mouse models. JCM and TJH designed and supervised the study. CA and JCM wrote the manuscript with input from all authors.

## Acknowledgements

We thank Odessa Van Goethem and Veronique Benne for excellent technical support and maintenance of the mouse colonies. We thank Tom Misteli (NCI/NIH, USA), Michael Dewaele (VIB, Belgium), Paulo P. Amaral (Cambridge University, UK) and Andrew Blackford (Oxford University, UK) for helpful discussions throughout the project. CA was supported by an IWT fellowship (no. 141372) from the Belgian Research Foundation - Flanders (FWO) and a Gustave Boël - Sofina travel grant (V433017N) from the Koning Boudewijn Stichting, Belgium. This work was supported by JSPS KAKENHI Grant Number 17H03604 and MEXT KAKENHI Grant Number 26113005 granted to S.N. We thank the members of the Research Resource Center at the RIKEN Brain Science Institute for the generation of the PAS KO mice. Work in the T.H.J. laboratory was supported by the ERC (grant 339953). We thank Manfred Schmid for computational support.

---

**Figure S1:**
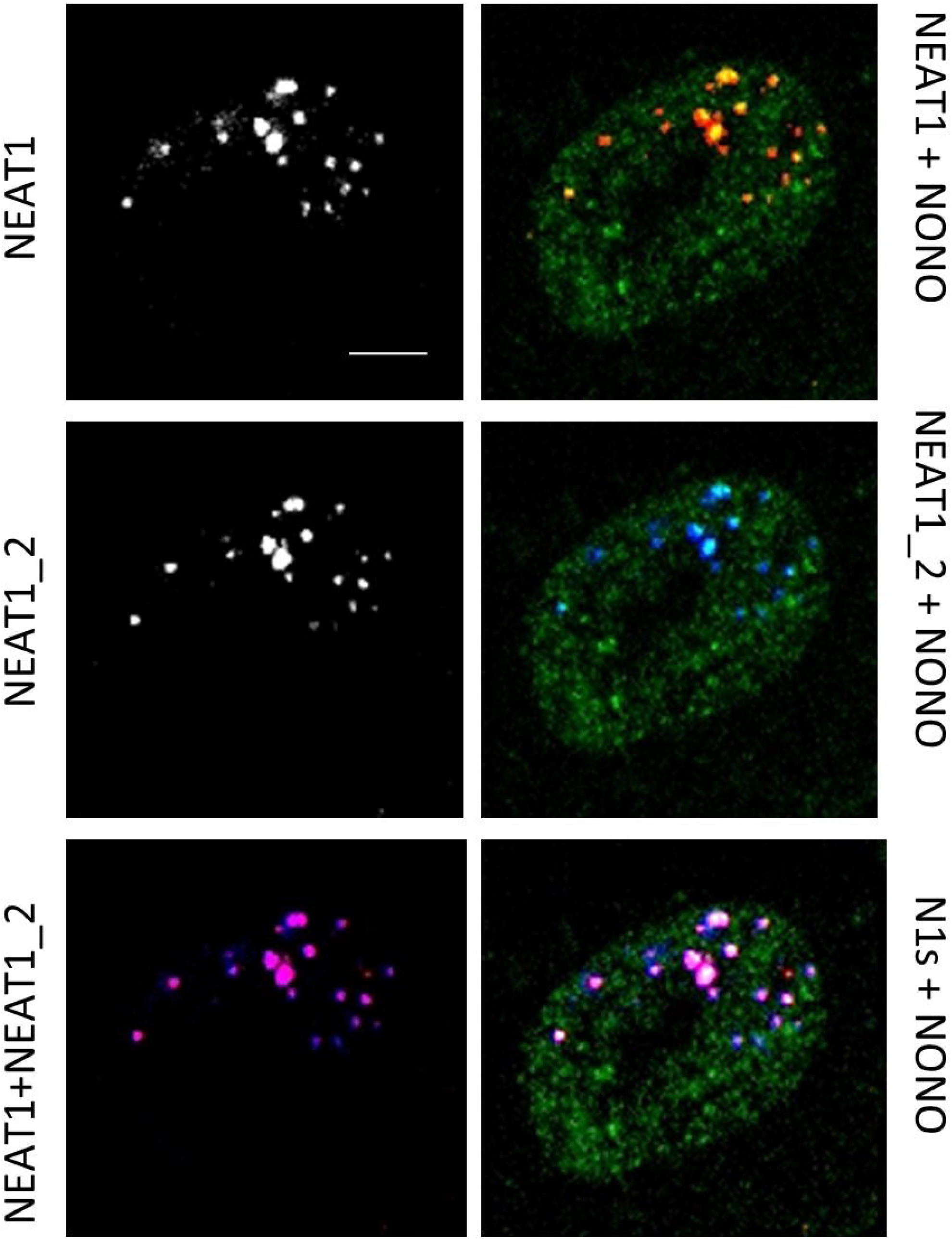
HU-induced *NEAT1_2* expression leads to PS formation. *NEAT1/NEAT1_2* RNA FISH of a HU treated cell (48h, 1 mM) combined with regular immunofluorescence against NONO, a canonical paraspeckle protein, showing paraspeckle integrity is preserved in HU conditions. Scale bar, 10 µm.

**Figure S2:**
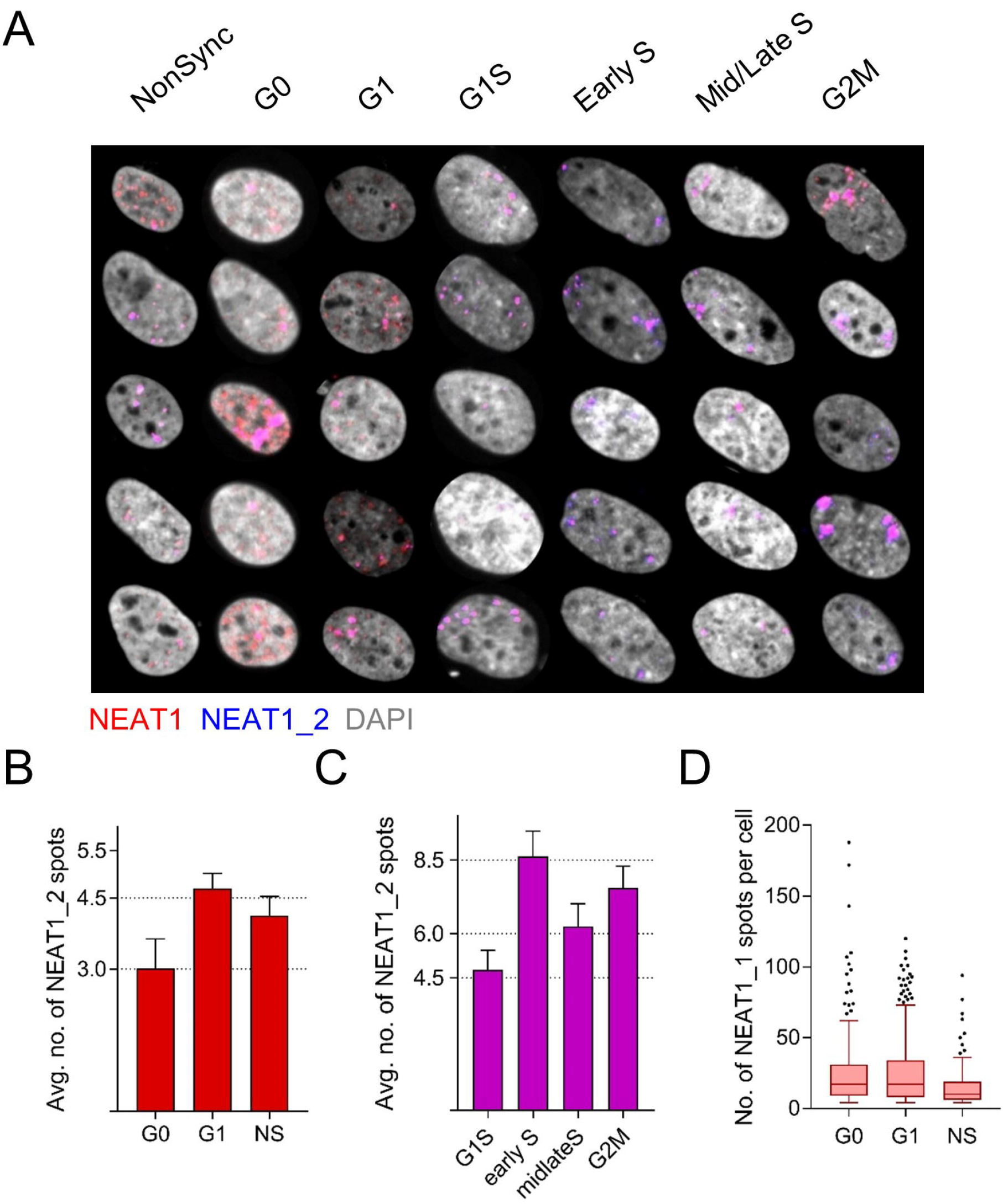
Monitoring PS formation and *NEAT1_1* levels in different cell cycle phases. A. Additional representative images showing the variation in RNA-FISH patterns of cells in the different cell cycle stages described in figure 2. For visualization purposes, cell sizes in this picture are not to scale. B-C. Quantification of the average number of *NEAT1_2* spots in cells in which *NEAT1_1* was detected (B) or not (C) in the majority of the cells. Bars show mean + standard error of individually quantified cells. D. Number of *NEAT1_1* speckles per cell in NS, G0 and G1 cells, Tukey boxplot representing one data point per cell.

**Figure S3:**
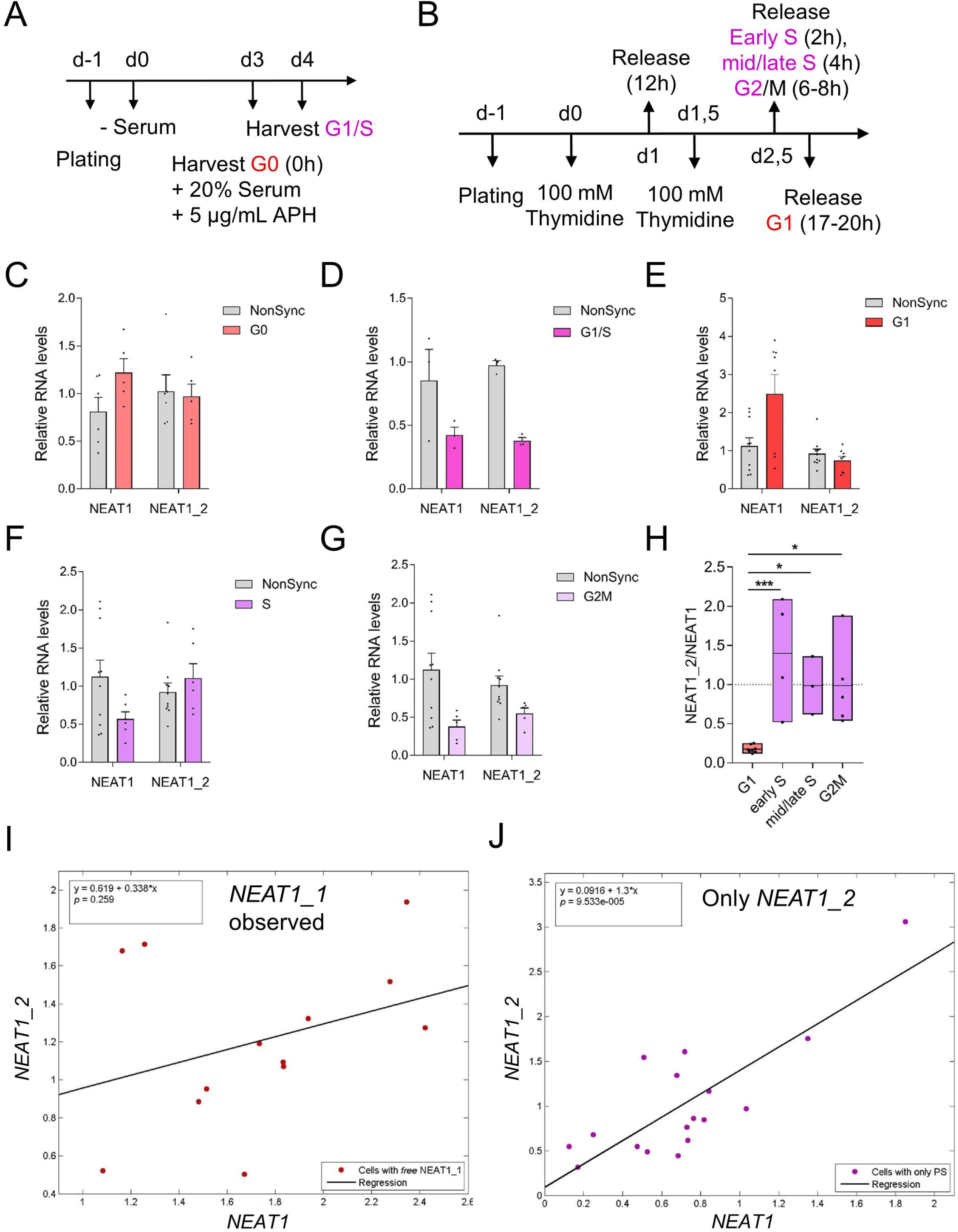
Monitoring expression of *NEAT1* isoforms in different phases of the cell cycle. A. Experiment layout for Figure 2 A-C. B. Experiment layout for Figure 2 D-E. C-G. Relative levels of *NEAT1* (both) and *NEAT1_2* isoforms by RT-qPCR analysis in G0 (C), G1/S (D), G1 (E), S (F) and G2M (G) phases of the cell cycle as synchronized in A & B of this figure. Each data point is an independent experiment. Bars and error depict mean + sem. H. *NEAT1_2* long isoform levels relative to the total of *NEAT1* detected in the cells in the experiments in figure 2 D-F. Bars are min to max with each data point an independent experiment. Lines depict the mean. Significance was tested by 1-way ANOVA with Sidak correction for multiple comparisons. *, p<0.05, ***, p<0.001. I-J, Pearson’s correlation analysis of relative levels of NEAT1_2 over total levels of *NEAT1* in RT-qPCR corrected for primer efficiencies in cells with primarily *NEAT1_1* detected outside of paraspeckles (I, r^2^ = 0,1141) and cells with only *NEAT1_2* (“only PS”) detected (J, r^2^ = 0,6488) in the RNA-FISH analysis in figure 2 D-F.

**Figure S4:**
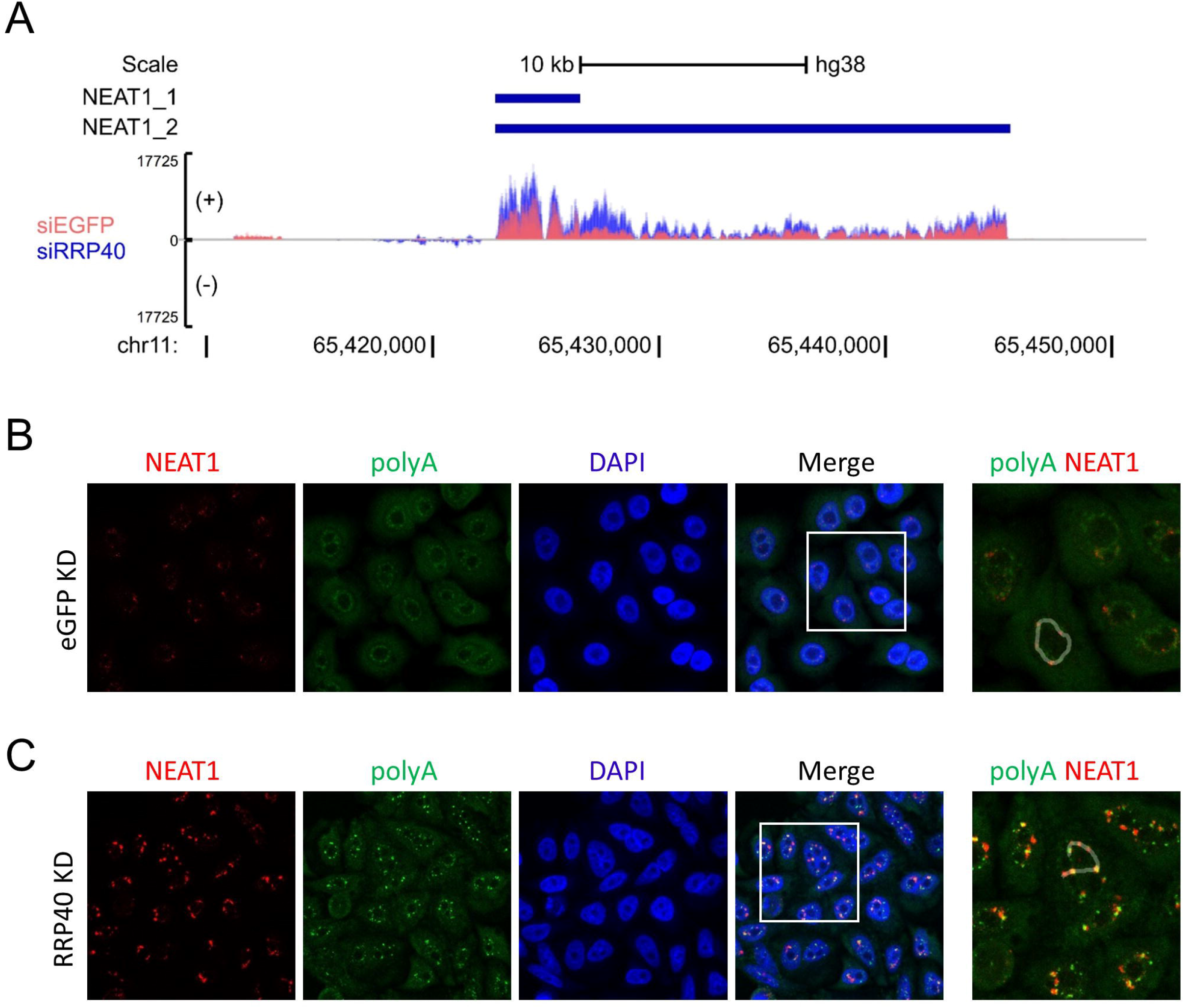
RRP40 KD causes *NEAT1_1* accumulation. A. RNA-seq tracks of the *NEAT1* locus in which RRP40 is knocked down. B. Image from which the line plot in figure 3D is derived. C. Image from which the line plot in figure 3E is derived. White lines represent lines along which the intensities of both channels are measured in Figure 3D&E.

**Figure S5:**
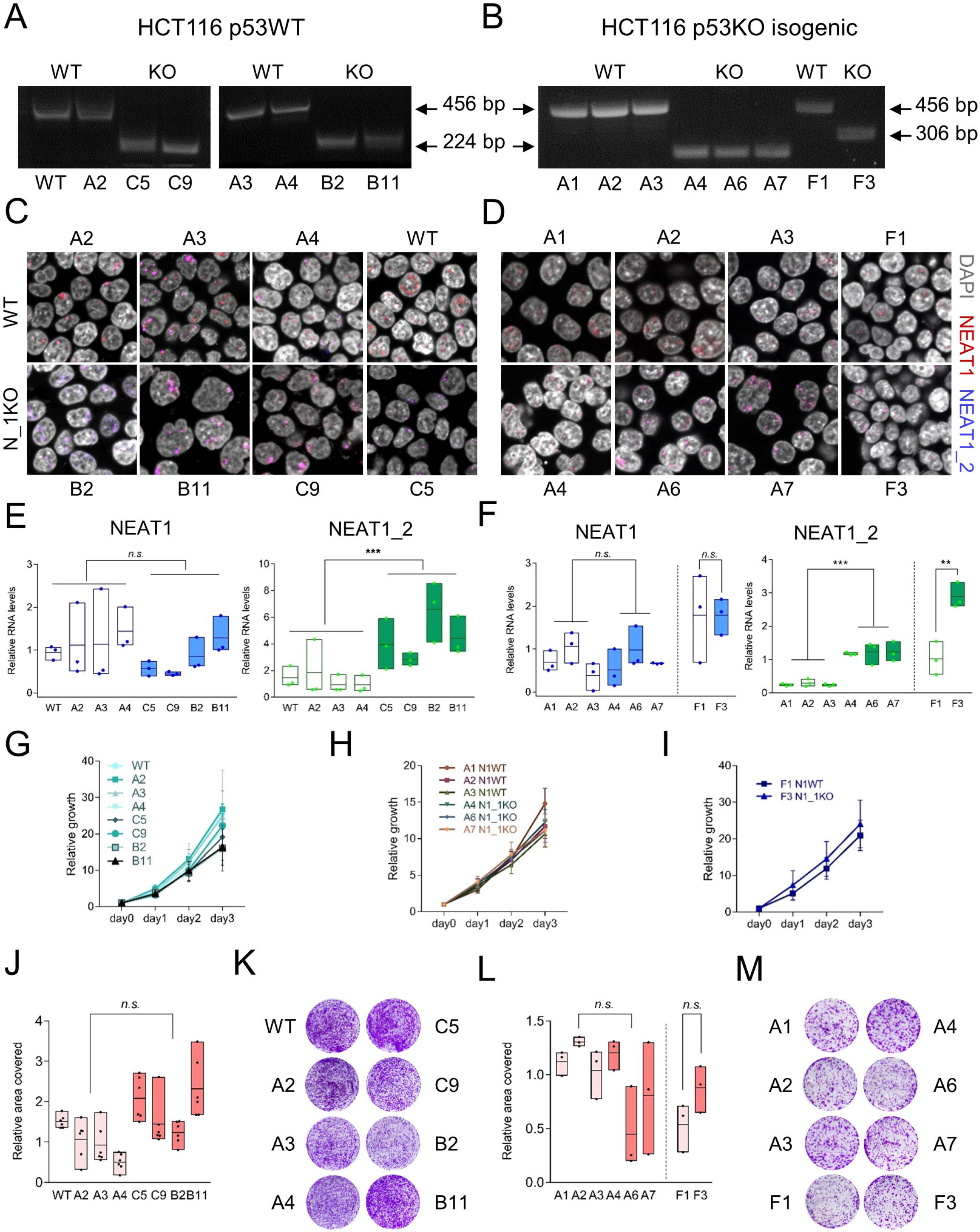
*NEAT1_1* KO does not induce HCT116 growth defects. A-B. Gel images of genotyping PCR using the primers in figure 4 for single-cell derived clones from HCT116 p53 wild type (A) and the isogenic p53 KO colon carcinoma cell line using 2 different guide RNA sets. C-D. Representative *NEAT1/NEAT1_2* RNA-FISH images for the HCT116 clones in A (C) and B (D). E-F. RT-qPCR analysis of NEAT1 total and *NEAT1_2* long isoforms. Dots represent individual data points of independent replicates. Statistical significance was calculated with 1-way ANOVA and Tukey’s correction for multiple testing with ***, p<0.001 and n.s., not significant. G-I: WST-1 analysis of short term growth in HCT116 WT (G), p53 KO gRNA set 1 (H) and set 2 (I) derived clones. J-M: long term growth analysis (2 weeks) of clones in A & B (J, L) and their representative well images after staining (K, M). Cells seeded in J-K: 6000. Cells seeded in L-M: 2000. Statistical significance was calculated with 1-way ANOVA and Tukey’s correction for multiple testing with n.s., not significant.

**Figure S6:**
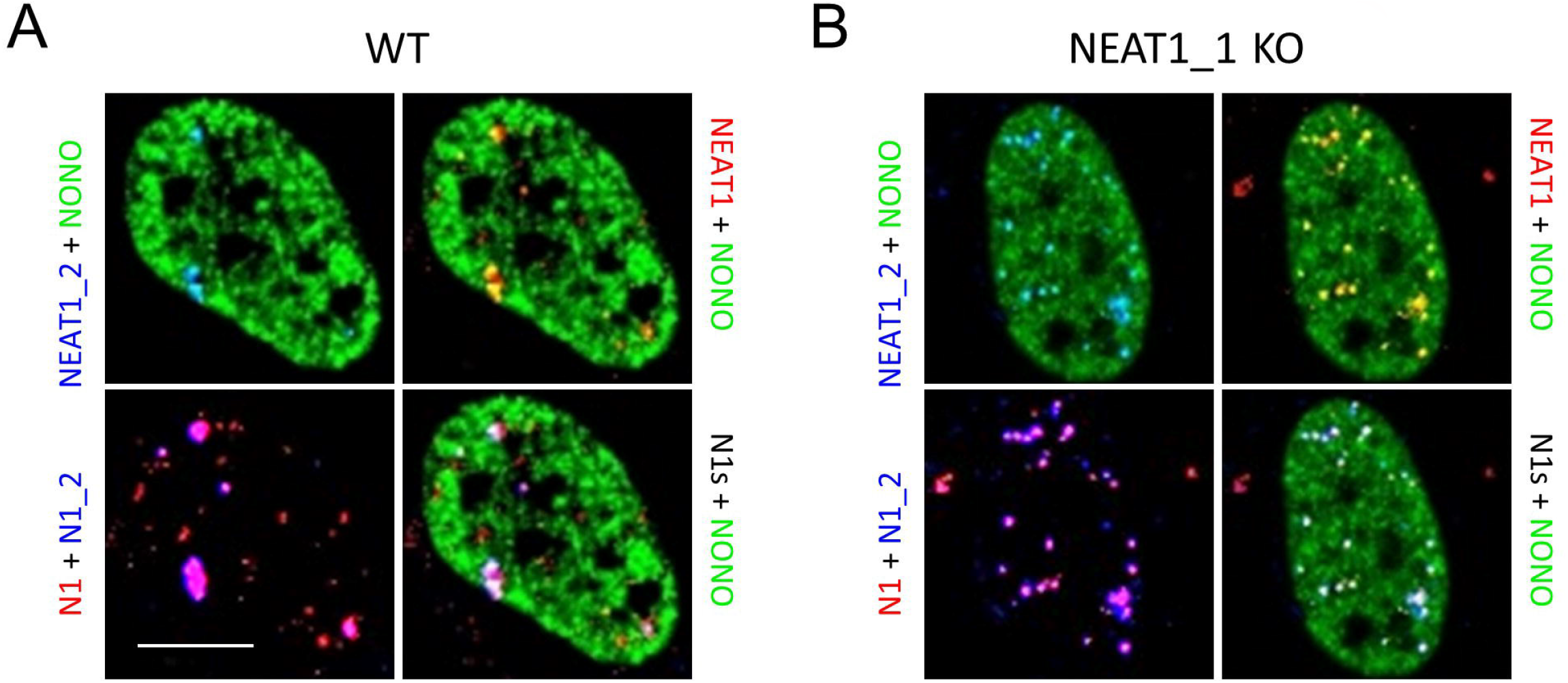
*NEAT1_1* KO cells make PS. A. *NEAT1/NEAT1_2* RNA FISH in wild type (WT) U2OS cells in G0 conditions in which the short isoform becomes prominently observed combined with IF for the paraspeckle protein NONO showing *NEAT1_2* but not *NEAT1_1* colocalization with the paraspeckles. B. Same as in A, but in *NEAT1_1* KO conditions lacking the short isoform. N1 = *NEAT1*; N1_2 = *NEAT1_2*. Scale bar, 10 um.

**Figure S7:**
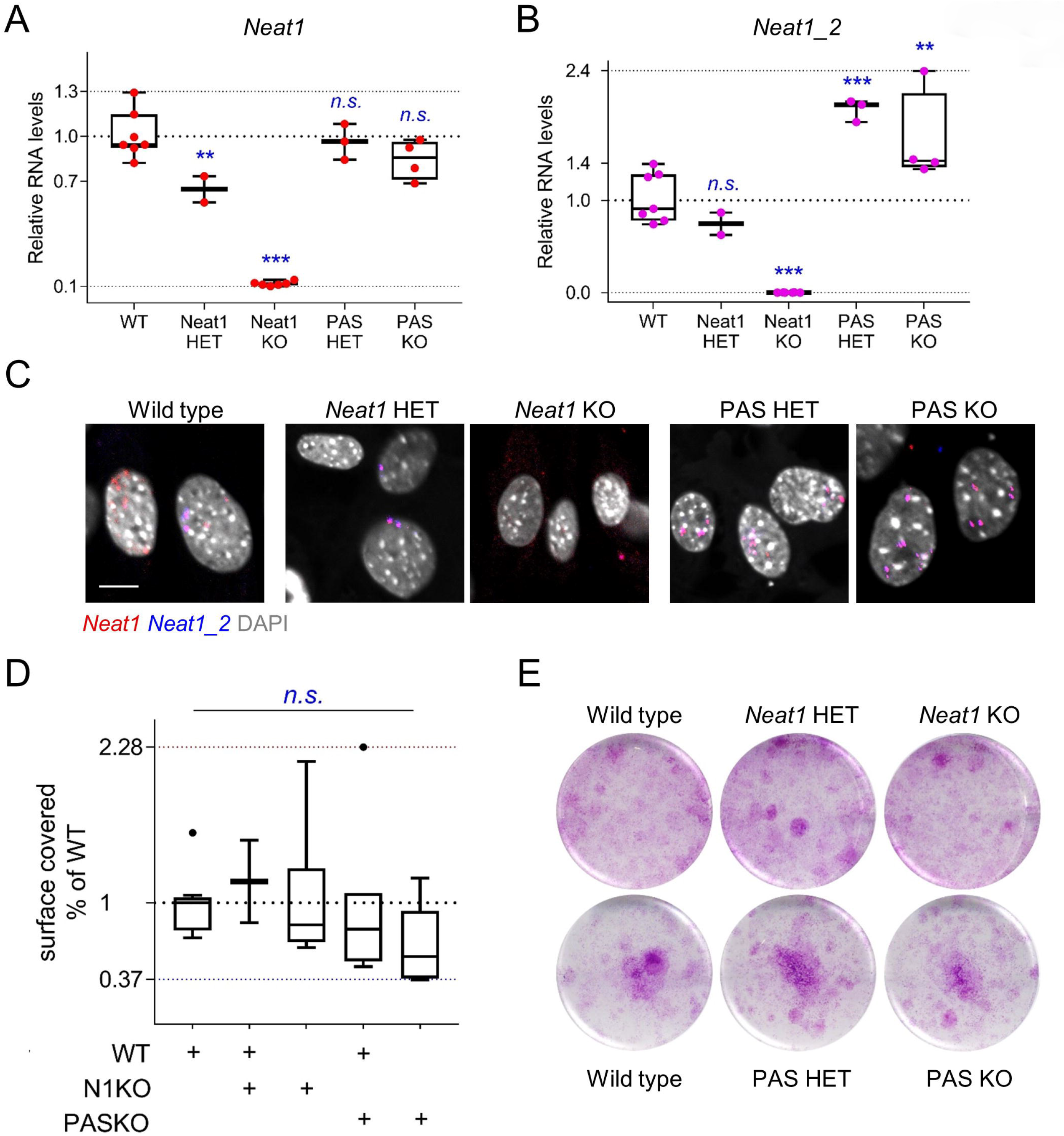
Loss of *Neat1_1* does not induce MEF growth defects. A-B. Min-to-max box plots of relative RNA levels by RT-qPCR for both *Neat1* isoforms (A), and the long *Neat1_2* isoform (B) in MEFs with the respective genotypes. C. Representative images from *Neat1/Neat1_2* RNA-FISH in these MEFs. Scale bar, 10 um. D-E. Tukey plots (D) and representative wells (E) of long term growth assay with 5000 cells seeded and stained after 2 weeks with *N* = at least 3 independently derived MEF clones per genotype. Each quantification is the average of 3 wells per MEF clone. For all graphs, significance was tested by 1-way ANOVA with Sidak correction for multiple comparisons. **, p<0.01, ***, p<0.001. n.s., not significant.

